# Murine GFP-Mx1 forms phase-separated nuclear condensates and associates with cytoplasmic intermediate filaments: novel antiviral activity against vesicular stomatitis virus

**DOI:** 10.1101/2020.08.12.248344

**Authors:** Pravin B. Sehgal, Hujuan Yuan, Mia F. Scott, Yan Deng, Feng-Xia Liang, Andrzej Mackiewicz

**Author notes:** Corresponding author: Dr. Pravin B. Sehgal, Rm. 201 Basic Science Building, Dept. Cell Biology & Anatomy, New York Medical College, Valhalla, NY 10595, USA. Huijuan Yuan, Division of Pulmonary, Allergy and Critical Care Medicine, Department of Medicine, University of Pittsburgh School of Medicine, Pittsburgh, PA 15213, USA.

## Abstract

Type I and III interferons (IFNs) induce expression of the “myxovirus resistance proteins” MxA in human cells and its ortholog Mx1 in murine cells. Human MxA forms *cytoplasmic* structures, some tethered to intermediate filaments. In contrast, murine Mx1 mainly forms *nuclear* bodies. Both HuMxA and MuMx1 are antiviral towards influenza A virus (FLUAV) (an orthomyxovirus). However, it has long been considered that HuMxA, but *not* MuMx1, was antiviral towards vesicular stomatitis virus (VSV) (a rhabdovirus). We previously reported that the cytoplasmic human GFP-MxA structures in Huh7 hepatoma cells were phase-separated membraneless organelles (MLOs) (“biomolecular condensates”). In the present study we investigated whether nuclear murine Mx1 structures might also represent phase-separated biomolecular condensates. The transient expression of murine GFP-Mx1 in Huh7 hepatoma and Mich-2H6 melanoma cells led to the appearance of Mx1 nuclear bodies. These GFP-MuMx1 nuclear bodies were rapidly disassembled by exposing cells to 1, 6-hexanediol (5% w/v), or to hypotonic buffer (40-50 mosM), consistent with properties of membraneless phase-separated condensates. FRAP assays revealed that the GFP-MuMx1 nuclear bodies upon photobleaching showed a slow partial recovery (mobile fraction: ∼18%) suggestive of a gel-like consistency. Surprisingly, expression of GFP-MuMx1 in Huh7 cells also led to the appearance of MuMx1 in a novel cytoplasmic giantin-based intermediate filament meshwork and in cytoplasmic structures in 20-30% of transiently transfected Huh7 cells. Remarkably, Huh7 cells with cytoplasmic murine GFP-MuMx1 filaments, but not those with only nuclear bodies, showed antiviral activity towards VSV. Thus, murine GFP-Mx1 nuclear bodies comprised phase-separated condensates. Unexpectedly, GFP-MuMx1 associated with cytoplasmic giantin-based intermediate filaments in a subset of Huh7 cells, and, such cells showed antiviral activity towards VSV.

## Introduction

Membraneless organelles (MLOs) in the cytoplasm and nucleus formed by liquid-liquid phase-separated (LLPS) biomolecular condensates are increasingly viewed as critical in regulating diverse cellular functions (1-9). These functions include cell differentiation, cell signaling, immune synapse function, nuclear transcription, RNA splicing and processing, mRNA storage and translation, virus replication and maturation, antiviral mechanisms, DNA sensing, synaptic transmission, protein turnover and mitosis (reviewed in 1-9). MLOs include the nucleolus, nucleoporin channels, nuclear speckles and paraspeckles, nuclear promyelocytic leukemia (PML) bodies, nuclear Cajal bodies, cytoplasmic processing (P) bodies, germinal bodies, Balbiani bodies, Negri bodies, stress granules, translation promoting TIS granules and several more recent discoveries such as condensates of synapsin, of the DNA sensor protein cyclic GMP-AMP synthase (cGAS) in the cytoplasm, and of active transcription-associated condensates in the nucleus (1-9). Overall, these condensates have liquid-like internal properties [as investigated using fluorescence recovery after photobleaching (FRAP) assays] and are metastable changing to a gel or to filaments commensurate with the cytoplasmic environment (temperature, ionic conditions, physical deformation or cytoplasmic “crowding”), and the incorporation of additional proteins, RNA or DNA molecules or posttranslational modifications (1-9). Indeed, specific DNA and RNA molecules participate in the assembly of many such cytoplasmic and nuclear condensates, and in their functions (1-9).

There is also an extensive literature on the changes observed in phase-separated stress granules and P-bodies in diverse virus-infected cells (9-14). In parallel with these insights, it has now been recognized that replication and maturation of negative-strand RNA viruses [e.g. vesicular stomatitis (VSV), rabies (as in Negri bodies), Ebola and measles viruses] occurs in cytoplasmic phase-separated condensates typically involving the viral nucleocapsid (N) protein with or without additional viral proteins (15-21). Even the replication of a DNA virus such as Epstein-Barr virus involves phase-separated nuclear bodies (22).

We recently reported that the human interferon (IFN)-inducible “myxovirus resistance protein A” (MxA), which displays antiviral activity against several different classes of RNA- and DNA-containing viruses (23-25), exists in the cytoplasm in membraneless metastable condensates in structures which include respective viral nucleocapsid proteins (9, 26, 27), confirming a previous report from 2002 of MxA in membraneless cytoplasmic structures (28). Mx proteins are a family of large dynamin-like GTPases of molecular weight in the size range 60-70 kDa which readily oligomerize (23-25). While human MxA which has a cytoplasmic localization, the murine Mx1 ortholog predominantly localizes in “Mx domains” or bodies in the nucleus (23-25). Consistent with its nuclear localization, murine Mx1 has an antiviral activity against orthomyxoviruses (such as influenza A virus, FLUAV)) which have a nuclear step in their growth cycle, but is reported to be inactive against rhabdoviruses (such as vesicular stomatitis virus, VSV) which replicate exclusively in the cytoplasm (23-25, 29-31). In contrast, human MxA shows antiviral activity towards both FLUAV and VSV (23-25).

To clarify the nomenclature of the Mx protein family we adopt the gene lineage tracing presented by Busnadiego et al (32) as follows: most mammalian Mx proteins are formed from two distinct gene lineages (MxA or MxB) that arose from an ancient duplication event. Thus, humans have two Mx proteins – MxA and MxB (some investigators call these as human Mx1 and human Mx2 respectively). Although mice also have two Mx proteins – Mx1 and Mx2, both these are paralogous members of the human MxA lineage. The genuine ortholog of human MxB has been lost in rodent and felid lineages. Thus, human MxB (which is also called human Mx2) and murine Mx2 are *not* orthologous. In this article we use MxA or HuMxA for the human protein, and Mx1 or MuMx1 for the orthologous murine protein.

Human MxA forms disparate membraneless structures solely in the cytoplasm and has antiviral activity towards several RNA- and DNA-containing viruses including orthomyxo- and rhabdoviruses (23-29), while human MxB is mainly associated with the cytoplasmic side of nuclear pores and additional cytoplasmic membraneless structures and the full-length MxB has antiviral activity against HIV and other lentiviruses by blocking entry of viral components into the nucleus (33-37). Murine Mx1 is mainly in nuclear bodies (it has a C-terminal nuclear localization signal), while murine Mx2 is mainly in cytoplasmic structures (23-25, 29-31). In terms of antiviral activity, reports in the literature indicate that the nuclear murine Mx1 has antiviral activity towards influenza virus (FLUAV) but not vesicular stomatitis virus (VSV), while the cytoplasmic “granular” murine Mx2 has antiviral activity towards both FLUAV and VSV (23-25, 29-31). Parenthetically, rats have three Mx proteins Mx1, Mx2 and Mx3 (29, 30, 32, 38). Rat Mx1 and rat Mx2 are orthologs of human MxA, and are nuclear and cytoplasmic respectively (29-31, 38). However, the nuclear-predominant rat Mx1 has antiviral activity towards both FLUAV and VSV, while the cytoplasmic-predominant rat Mx2 is antiviral towards VSV only (29-31, 38). Rat Mx3, which is mainly dispersed in the cytoplasm, has little apparent antiviral activity (29-31, 38).

Engelhardt and colleagues (39, 40) observed that murine Mx1 nuclear bodies sometimes overlapped with or were in juxtaposition to PML bodies, nuclear speckles and Cajal bodies. The latter three structures have all now been identified as phase-separated biomolecular condensates involved in RNA handling (1-9, 41). Curiously, there was no obligatory relationship between nuclear Mx1 bodies and PML bodies in that the formation of Mx1 bodies was not altered in *PML*^*-/-*^ cells (40).

In the present study we investigated whether nuclear murine GFP-Mx1 bodies might also consist of phase-separated biomolecular condensates. The data obtained confirmed that murine GFP-Mx1 nuclear bodies had the properties of phase-separated biomolecular condensates in the nuclear compartment. Unexpectedly, we discovered that murine GFP-Mx1 was also associated with novel giantin-based intermediate filaments and with cytoplasmic bodies in Huh7 hepatoma cells. Remarkably, such cells expressing murine GFP-Mx1 in the cytoplasm, but not those containing only GFP-MuMx1 nuclear bodies, displayed a strong antiviral phenotype against the rhabdovirus VSV.

## Materials and Methods

### Cells and cell culture

Human hepatoma cell line Huh7 (42) was a gift from Dr. Charles M. Rice, The Rockefeller University. Human Mich2-H6 melanoma cells were developed by Mackiewicz and colleagues (43). The respective cell lines were grown in DMEM supplemented with 10% v/v fetal bovine serum (FBS) in T25 flasks (26, 27, 44, 45). For experiments, the Huh7 and Mich2-H6 cells were typically plated in 35 mm dishes without or with cover-slip bottoms (26, 27).

### Plasmids and transient transfection

The GFP (1-248)-tagged full-length human MxA and the GFP-tagged full-length murine Mx1 expression vectors were gifts of Dr. Jovan Pavlovic (University of Zurich, Switzerland) (46, 47); the GFP tag was located on the N-terminal side of the Mx coding sequence. Plasmid vectors for expression of the N1-GFP tag only and one for expression of GFP-STAT3 were used as negative controls (48, 49). Transient transfections were carried out using just subconfluent cultures in 35 mm plates using DNA in the range of 0.3-2 µg/culture and the Polyfect reagent (Qiagen, Germantown, MD) and the manufacturer’s protocol (with 10 µl Polyfect reagent per 35 mm plate).

### VSV stock and virus infection

A stock of the wild-type Orsay strain of VSV (titer: 9 × 10^8^ pfu/ml) was a gift of Dr. Douglas S. Lyles (Dept. of Biochemistry, Wake Forest School of Medicine, Winston-Salem, NC). Virus infection was carried out essentially as described by Carey et al (50) as summarized in Davis et al using Huh7 cells (27). Briefly, Huh7 cultures (approx. 2 × 10^5^ cells per 35 mm plate), previously transfected with the pGFP-Mx1 expression vector (1-2 days earlier), were replenished with 0.25 ml serum-free Eagle’s medium and 10-20 µl of the concentrated VSV stock added (corresponding to MOI >10 pfu/cell). The plates were rocked every 15 min for 1 hr followed by addition of 1 ml of full culture medium. For the experiment shown in Fig. 9, the cultures were fixed at 4 hr after the start of the VSV infection and immunostained for VSV nucleocapsid (N) protein using an mAb provided by Dr. Douglas S. Lyles (mAb 10G4). N-protein immunofluorescence (in red) in GFP-positive (nuclear or cytoplasmic) and negative cells was quantitated on a per cell basis as summarized in Davis et al (27) and expressed in arbitrary units (AU) as integrated intensity/cell.

**Fig. 1.**
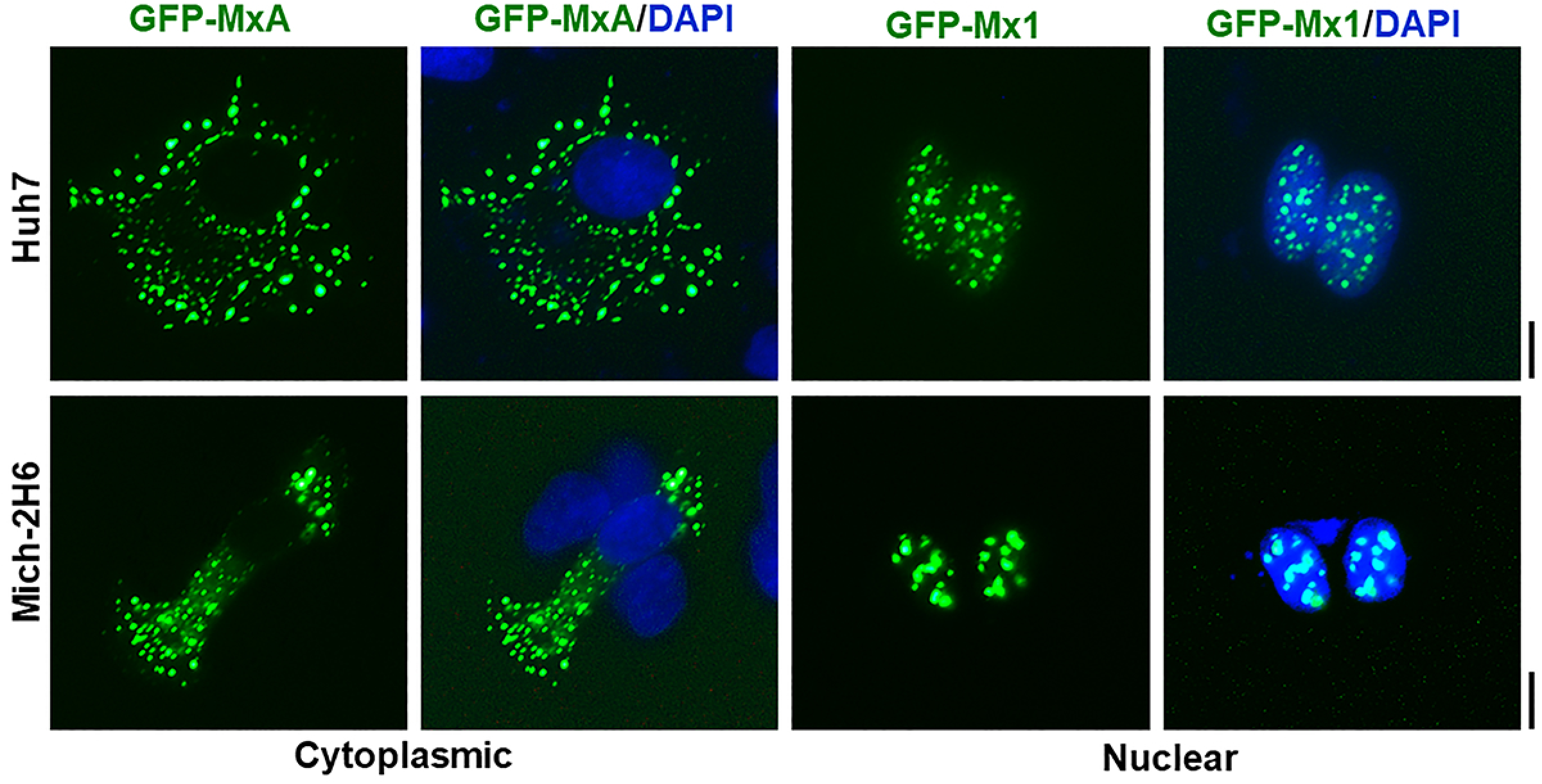
Comparison of subcellular localization of human GFP-MxA with murine GFP-Mx1 in two different cell lines. Cultures of Huh7 and Mich-2H6 cells in 35 mm plates were transiently transfected with expression vectors for human GFP-MxA or murine GFP-Mx1, fixed using 4% paraformaldehyde 2 days later, additionally stained with DAPI to visualize the nuclei, and imaged using two-color fluorescence. The figure illustrates representative cells. None of the cells transfected with human GFP-MxA vector showed any MxA in nuclei; 70-80% of Huh7 cells transfected with GFP-Mx1 showed only nuclear Mx1 (for cells with cytoplasmic GFP-Mx1 structures see Figs. 6-8); almost all of Mich-2H6 cells transfected with GFP-Mx1 showed only nuclear GFP-Mx1 bodies. Scale bar = 10 µm.

**Fig. 2.**
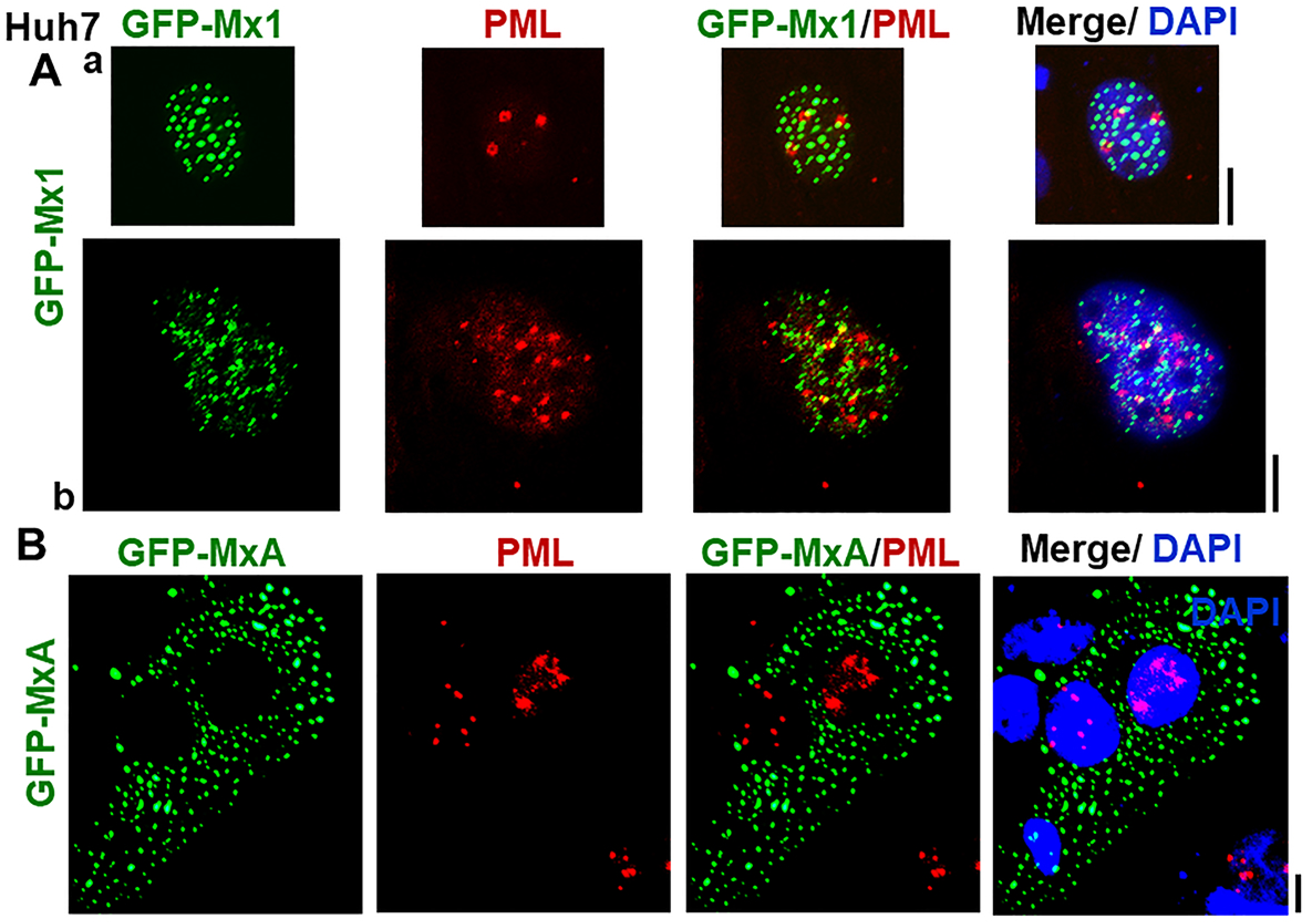
Lack of colocalization of nuclear murine GFP-Mx1 with nuclear PML bodies. Cultures of Huh7 cells in 35 mm plates were transiently transfected with expression vectors for murine GFP-Mx1 (Panel A) or human GFP-MxA (Panel B), fixed using 4% paraformaldehyde 2 days later, probed for PML using immunofluorescence methods, additionally stained with DAPI to visualize the nuclei, and imaged using three-color fluorescence. The figure illustrates representative cells. Pearson’s correlation coefficients R (after Costes’ automatic thresholding) in Panels Aa, Ab and B were 0.0, 0.25 and 0.0 respectively. Scale bars = 10 µm.

**Fig. 3.**
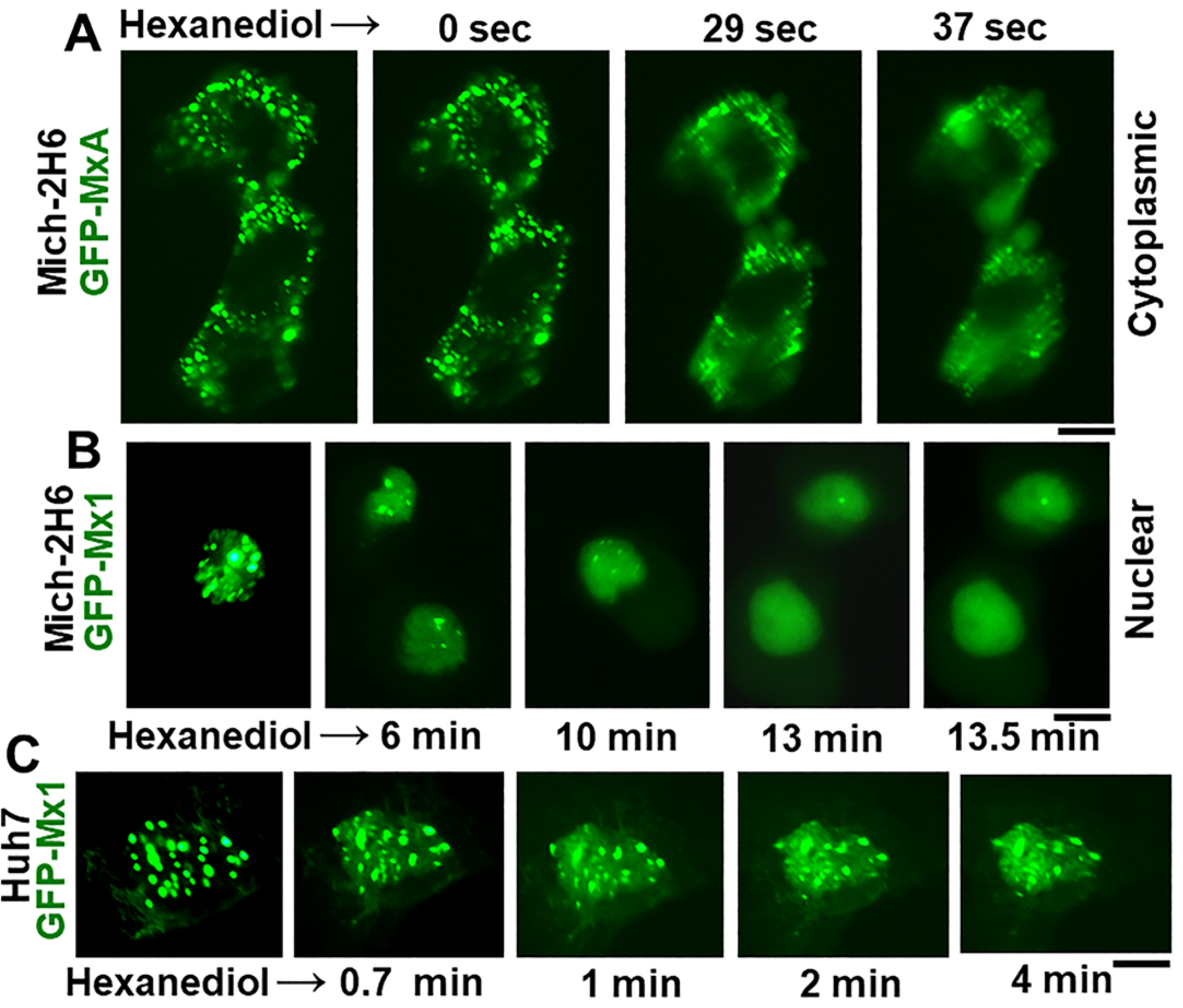
Disassembly of nuclear GFP-Mx1 bodies by 1,6-hexanediol. Huh7 and Mich-2H6 cells were transfected with either GFP-MxA vector (as a positive control) or GFP-Mx1 vector. One-two days later the live cells were first imaged in PBS, and then the culture medium was changed to PBS supplemented with 5% (w/v) 1,6-hexanediol followed by imaging at different times thereafter (9, 27). Panel A, illustrates a positive control verifying the rapid disassembly of cytoplasmic GFP-MxA condensates (9, 27) in Mich-2H6 cells. Panels B and C illustrate a somewhat slower disassembly of GFP-Mx1 nuclear bodies after hexanediol exposure in both Mich-2H6 and Huh7 cells. Scale bar = 10 µm.

**Fig. 4.**
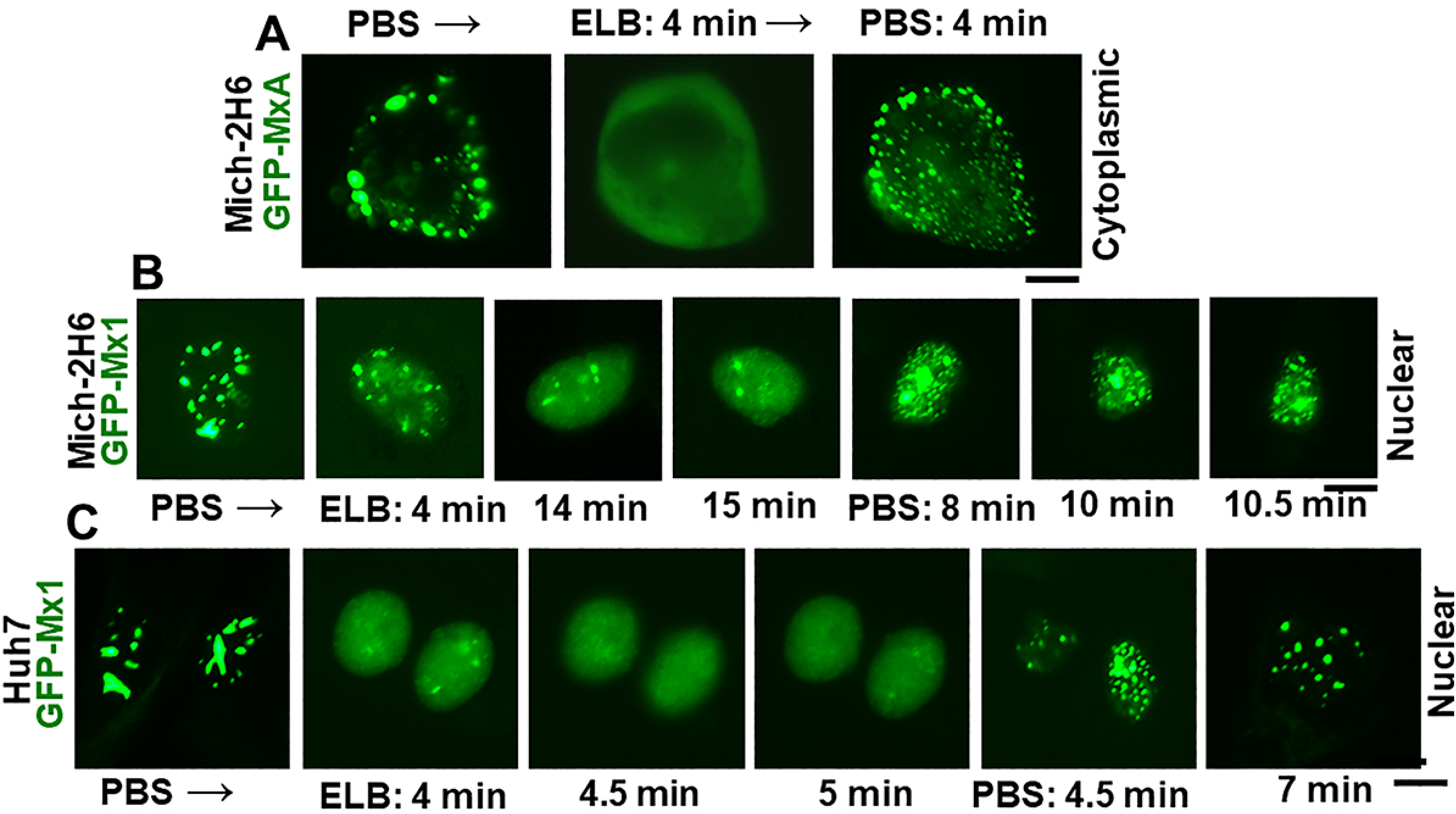
Hypotonic disassembly and isotonic reassembly of nuclear GFP-Mx1 bodies. Mich-2H6 or Huh7 cells were transiently transfected either with the GFP-MxA vector (as a positive control) or the GFP-Mx1 vector. One-two days later the live cells were first imaged in PBS, and then the culture medium was changed to hypotonic ELB buffer (40 mOsm) followed by imaging at different times thereafter (9, 27). After 10-15 min, the culture medium was changed to isotonic PBS and live cell imaging continued for another 10-15 min. The figure illustrates representative cells at different time points. Panel A, illustrates a positive control verifying the rapid disassembly of cytoplasmic GFP-MxA condensates (9, 27) in Mich-2H6 cells upon hypotonic exposure and reassembly of the condensates upon shifting the culture medium to isotonic PBS. A time-lapse movie of this experiment is shown in Supplemental Movie S1. Panels B and C illustrate a somewhat slower disassembly of GFP-Mx1 nuclear bodies after hypotonic exposure and isotonic reversal in both Mich-2H6 and Huh7 cells. Scale bar = 10 µm.

**Fig. 5.**
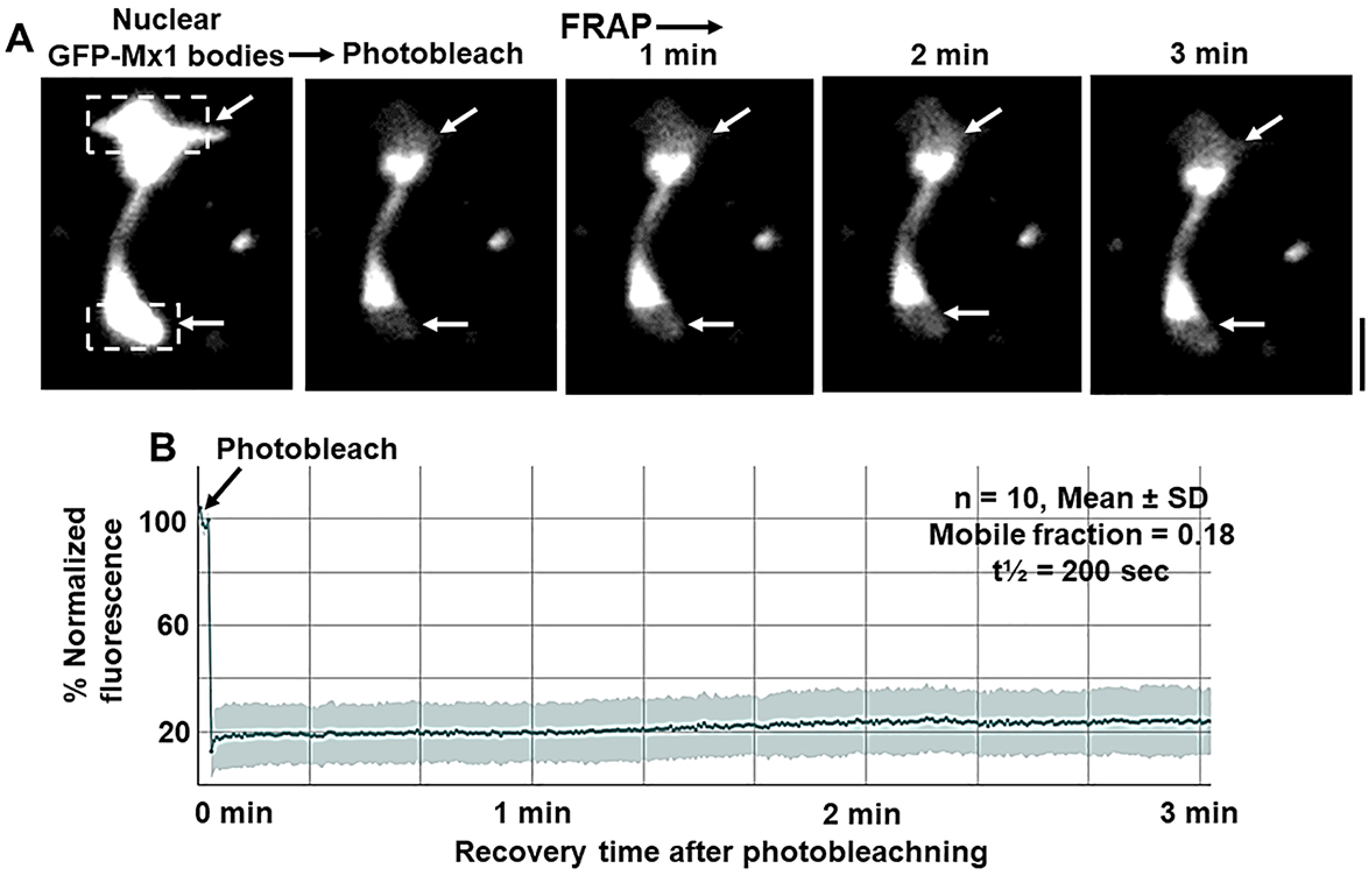
FRAP analyses of GFP-Mx1 in nuclear bodies. Huh7 cells grown in 35 mm coverslip-bottom plates (MatTek) were transiently transfected with the GFP-Mx1 vector and cells imaged 2 days later using Zeiss 880 confocal microscopy system. Nuclear bodies in selected cells were subjected to photobleaching (dashed rectangles) and the recovery from bleaching monitored for 3 minutes (Panel A); scale bar = 5 um. Supplemental Movie S2 is the time-lapse movie corresponding to the images in Panel A. Panel B summarizes numerical data derived from ten such experiments.

**Fig. 6.**
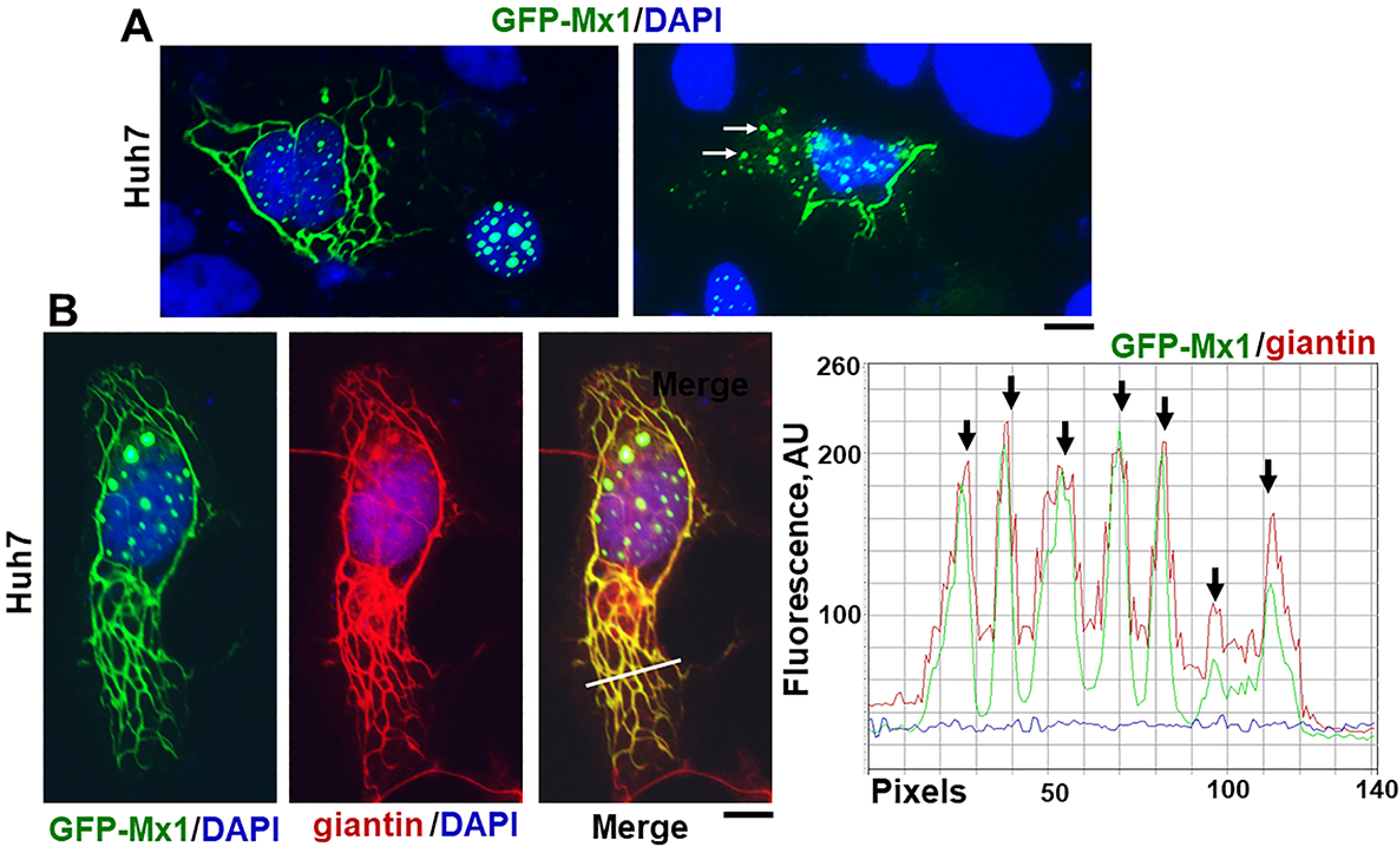
Murine GFP-Mx1 localized to cytoplasmic structures in human Huh7 cells. Cultures of Huh7 cells in 35 mm plates were transiently transfected with the expression vector for murine GFP-Mx1. Two days later the cultures were fixed using 4% paraformaldehyde, permeabilized using the 0.1% Triton buffer, and stained with DAPI to visualize the nuclei, and imaged using two-color fluorescence. In Panel A, the left image shows two adjacent cells – one showing GFP-Mx1 exclusively in nuclear bodies and one cell with GFP-Mx1 present extensively in cytoplasmic filaments and cytoplasmic structures. Similarly, the right image in panel A shows adjacent cells – one with extensive cytoplasmic filaments and cytoplasmic bodies of GFP-Mx1 And part of a nucleus with only nuclear GFP-Mx1 bodies (at lower left). Approximately 20-30% of transfected cells showed cytoplasmic murine GFP-Mx1. Panel B summarizes a three-color immunofluorescence analysis of the cytoplasmic GFP-Mx1 filamentous structures for intermediate filaments (using the anti-giantin pAb). Red and green pixel fluorescence was measured along the white line in the merged image in Panel B and the data summarized in the plot next to the right of the image. Arrows in the plot indicate colocalization of red and green filaments. Scale bars = 10 µm.

**Fig. 7.**
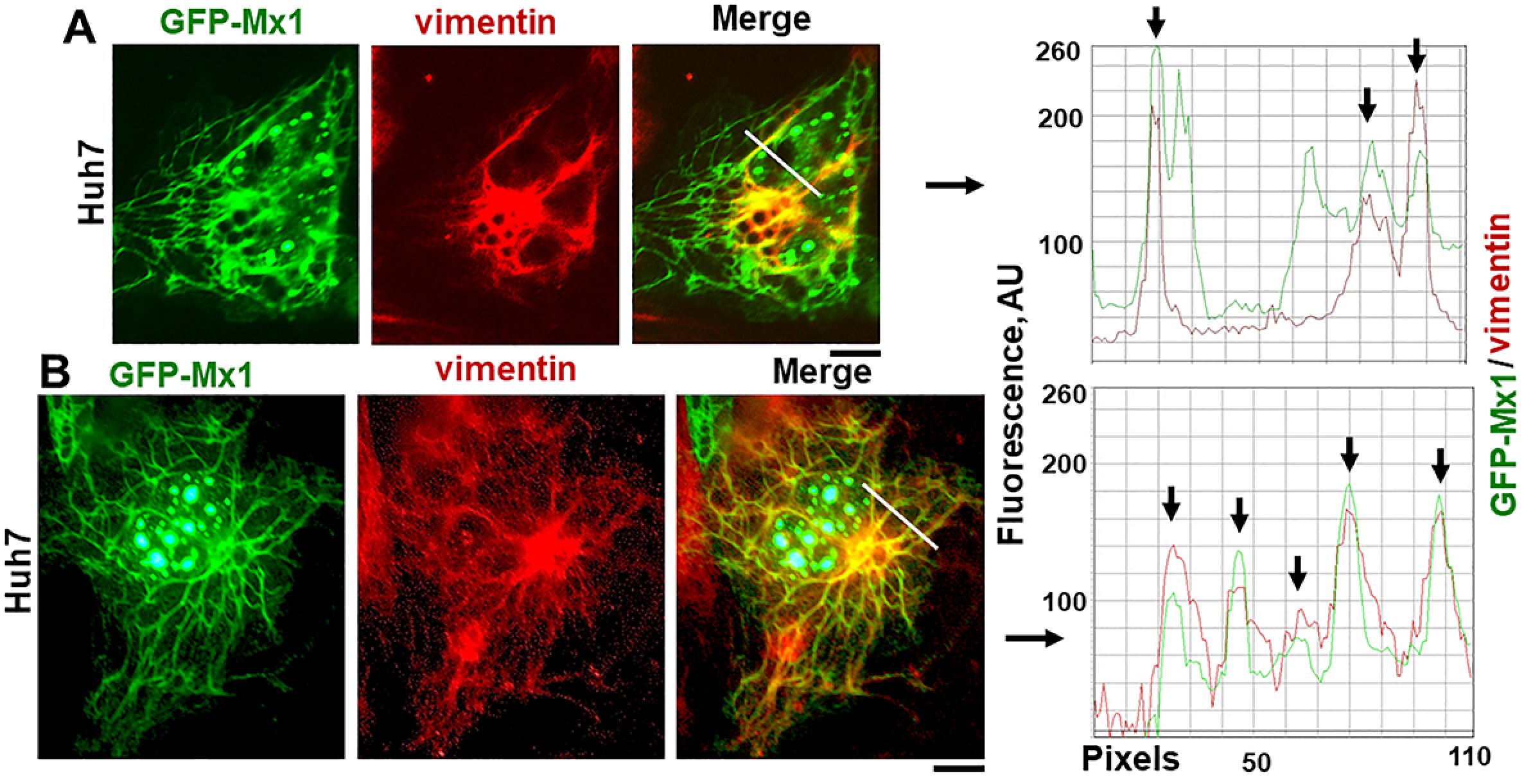
Murine GFP-Mx1 localized to intermediate filaments in human Huh7 cells. Cultures of Huh7 cells in 35 mm plates were transiently transfected with the expression vector for murine GFP-Mx1. Two days later the cultures were fixed using 4% paraformaldehyde, permeabilized using the 0.1% Triton buffer, immunostained with anti-vimentin mAb to visualize intermediate filaments and then with stained with DAPI to visualize the nuclei. The cultures were imaged using three-color fluorescence. Panels A and B show two examples of cells with GFP-Mx1 in cytoplasmic filamentous structures some of which also overlap vimentin filaments. Fluorescence in red and green pixels along the white lines in the merged images in panels A and B are plotted to the right of the respective images. Arrows indicate colocalized filamentous structures (GFP and vimentin). Scale bars = 10 µm.

**Fig. 8.**
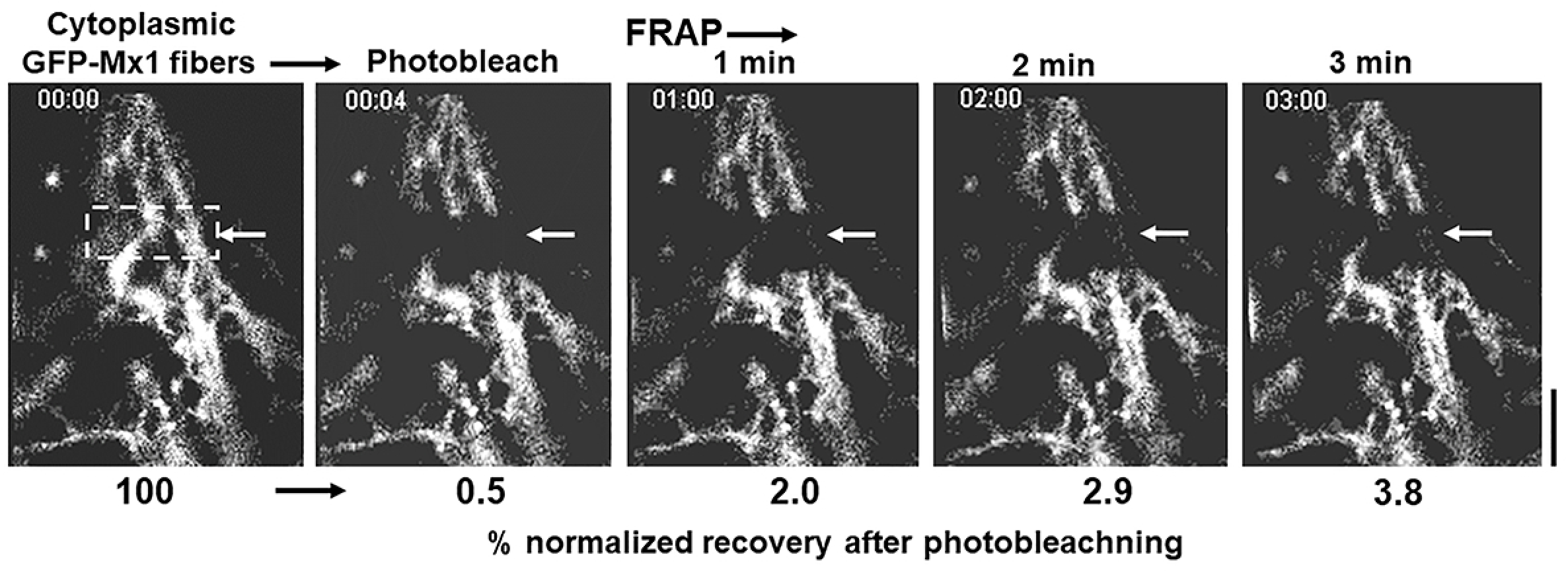
FRAP analyses of GFP-Mx1 in cytoplasmic filamentous meshwork. Huh7 cells grown in 35 mm coverslip-bottom plates (MatTek) were transiently transfected with the GFP-Mx1 vector and cells imaged 2 days later using Zeiss 880 confocal microscopy system. A rectangular section of (dashed lines) of a cell with GFP-Mx1 in cytoplasmic filaments was subjected to photobleaching and the recovery from bleaching monitored for 3 minutes; scale bar = 20 um. The numerals below each panel summarize the % normalized recovery after photobleaching. Supplemental Movie S3 is the time-lapse movie corresponding to this experiment.

**Fig. 9.**
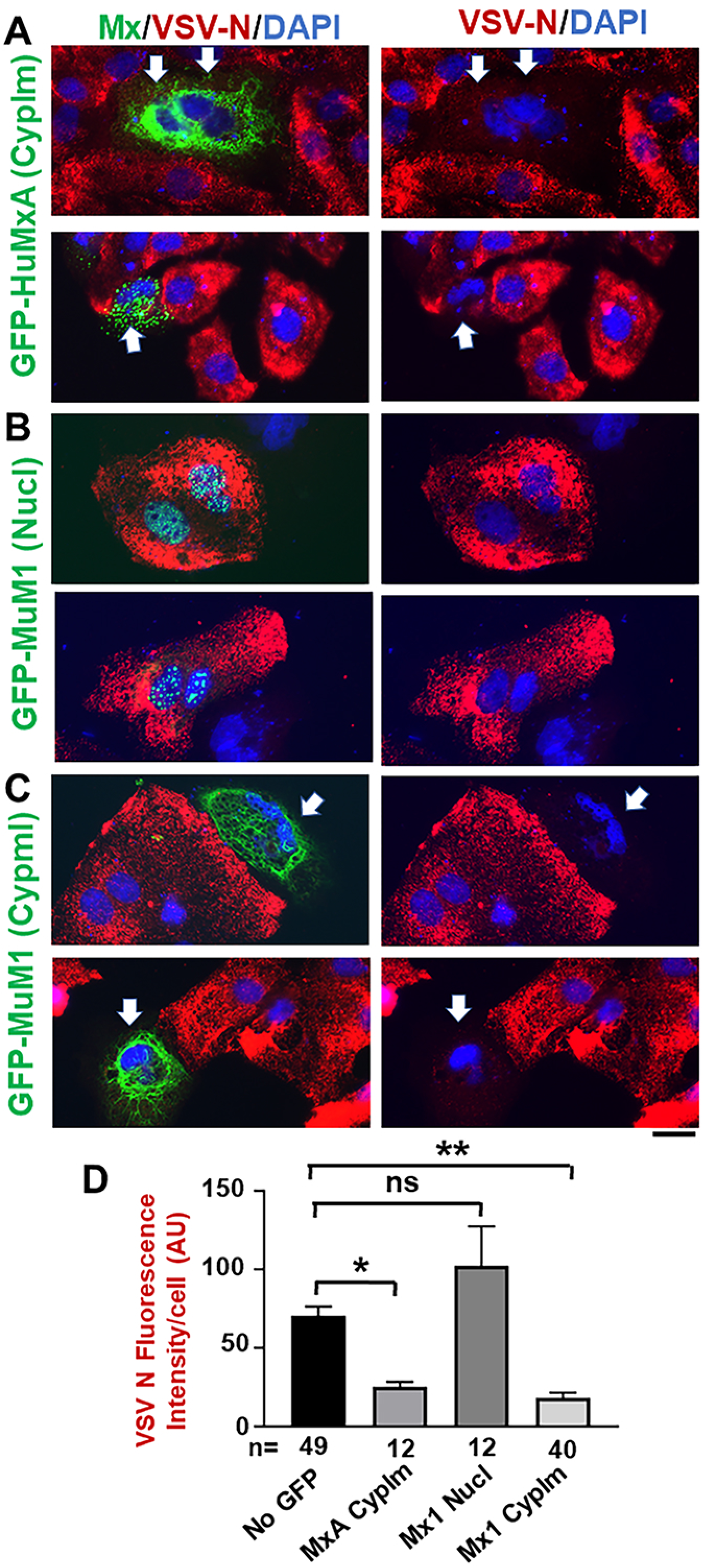
Antiviral phenotype of Huh7 cells with nuclear or cytoplasmic GFP-MuMx1 towards VSV. Huh7 cells (approx. 2 × 10^5^) per 35 mm plate, transfected with the GFP-HuMxA or GFP-MuMx1 expression vectors 2 days earlier, were replenished with 0.25 ml serum-free Eagle’s medium and then 20 µl of a concentrated VSV stock of the wt Orsay strain added (corresponding to multiplicity of infection >10 plaque forming units/cell). The plates were rocked every 15 min for 1 hr followed by addition of 1 ml of full culture medium. The cultures were fixed with 4% paraformaldehyde at 4 hr after the start of the VSV infection and the extent of VSV N protein expression in individual cells evaluated using immunofluorescence methods (using the mouse anti-N mAb) and Image J for quantitation (27). Panels A, B and C illustrate representative cells showing the absence of any GFP, or the appearance of cytoplasmic GFP-HuMxA, nuclear GFP-MuMx1 or cytoplasmic GFP-MuMx1 and the corresponding level of expression of viral N protein (thick arrows point to cells displaying an antiviral effect). All scale bars = 20 μm. Panel D, enumerates N protein expression in various classes of cells shown in Panels A, B and C imaged at identical exposure settings and expressed in arbitrary fluorescence units (AU) per cell; mean ± SE, n= number of cells evaluated per group in this experiment. Statistical significance was evaluated using ANOVA (Kruskal-Wallis with Dunn’s post-test for multiple comparisons); * *P* < 0.01; ** *P* <0.001; ns, not significant (*P* > 0.05).

### Live-cell fluorescence imaging

Live-cell imaging of GFP-MxA and GFP-Mx1 structures in transiently transfected cells was carried out in cells grown in 35 mm plates using the upright the Zeiss AxioImager 2 equipped with a warm (37°C) stage and a 40x water immersion objective, and also by placing a coverslip on the sheet of live cells and imaging using the 100x oil objective (as above) with data capture in a time-lapse or z-stack mode (using Axiovision 4.8.1 software)(27).

FRAP experiments were performed using a Zeiss LSM880 confocal microscope with Zeiss Plan-Apochromat 63x/1.4 Oil objective on randomly picked cells grown in cover-slip bottomed 35 mm culture dishes (27). Five pre-bleach images were acquired before 25 cycles of bleaching with full power of Argon 488nm laser on small area (∼3 μm^2)^ which reduces intensity to ∼50%. Fluorescence recovery was monitored continuously for 180 seconds at 2 frames/sec. All image series were aligned and registered with Fiji ImageJ’s SIFT plugin and FRAP analyses of MF and t1/2 were performed afterwards with a Jython script (https://imagej.net/Analyze_FRAP_movies_with_a_Jython_script) incorporated into Image J (Fiji).

### Phase transition experiments

Live GFP-Mx1 or GFP-MxA expressing Huh7 hepatoma cells in 35 mm plates were imaged using a 40x water-immersion objective 2-5 days after transient transfection in growth medium or serum-free DMEM medium or in phosphate-buffered saline (PBS). After collecting baseline images of Mx condensates (including time-lapse sequences), the cultures were exposed to 1,6-hexanediol (5% w/v) in PBS, or to hypotonic buffer (ELB; 10 mM NaCl, 10 mM Tris, pH 7.4, 3 mM MgCl_2_) and live-cell time-lapse imaging continued (9, 27). After approximately 5-10 min the cultures exposed to hypotonic ELB were replenished with isotonic phosphate-buffered saline (PBS) and imaged for another 5-10 min.

### Immunofluorescence imaging

Typically, the cultures were fixed using cold paraformaldehyde (4%) for 1 hr and then permeabilized using a buffer containing digitonin or saponin (50 µg/ml) and sucrose (0.3M) (27); cultures for immunostaining of nuclear bodies, vimentin or VSV N protein were permeabilized using 0.05% Triton in PBS. Single-label and double-label immunofluorescence assays were carried out using antibodies as indicated, with the double-label assays performed sequentially. Fluorescence was imaged as previously reported (26, 27, 44, 45, 49) using an erect Zeiss AxioImager M2 motorized microscopy system with Zeiss W N-Achroplan 40X/NA0.75 water immersion or Zeiss EC Plan-Neofluor 100X/NA1.3 oil objectives equipped with an high-resolution RGB HRc AxioCam camera and AxioVision 4.8.1 software in a 1388 × 1040 pixel high speed color capture mode. Images in z-stacks were captured using Zeiss AxioImager software; these stacks were then deconvolved and rendered in 3-D using the 64-bit version of the Zeiss AxioVision software. Deconvolution of 2-D images was carried out using Image J (Fiji) software. Colocalization analyses were carried out using Image J software (Fiji) and line scans were carried out using AxioVision 4.8.1 software. High-resolution immunofluorescence/ fluorescence imaging of selected cultures was carried out using a Zeiss Confocal 880 Airyscan system (100x oil/1.46-NA objective).

### Antibody reagents

Rabbit pAb to human MxA (H-285) (sc-50509), and mouse mAb vimentin (V9) (sc-6260) were purchased from Santa Cruz Biotechnology Inc. (Santa Cruz, CA). Rabbit pAb to giantin (1-469 fragment) (ab24586) was purchased from Abcam Inc. (Cambridge, MA); a different rabbit pAb to giantin (1-469 fragment) (Poly19243) was also purchased from BioLegend Inc. (San Diego, CA). Mouse mAb to the VSV nucleocapsid (N) designated 10G4 was a gift from Dr. Douglas S. Lyles (Wake Forest School of Medicine, NC). Blocking peptides corresponding to human STAT2 and human giantin (1-469) were purchased from Santa Cruz Biotechnology, Inc. or custom synthesized by GeneScript USA Inc. (Piscataway, NJ) respectively. Respective AlexaFluor 488- and AlexaFluor 594-tagged secondary donkey antibodies to rabbit (A-11008 and A-11012) or mouse (A-21202 and A-21203) IgG were from Invitrogen Molecular Probes (Eugene, OR).

### Statistical testing

This was carried out (as in Fig. 9D) using non-parametric one-way ANOVA (Kruskal-Wallis) with Dunn’s post-hoc test for multiple comparisons.

## Results

### Murine GFP-Mx1 forms nuclear bodies

In contrast to the transient expression of human GFP-MxA in Huh7 hepatoma and Mich-2H6 melanoma cells which led to formation of cytoplasmic condensates of MxA (Fig. 1, left half), the transient expression of murine GFP-Mx1, as expected (23-25), led to formation nuclear bodies in most cells (Fig. 1, right half). These data confirm the localization of these respective proteins as reported in the literature (23-25). Moreover, in confirmation of the observations of Engelhardt et al (40), the nuclear bodies formed by murine GFP-Mx1 in Huh7 cells were distinct from nuclear PML bodies (Fig. 2A). As a negative control, cytoplasmic GFP-MxA condensates also did not include PML (Fig. 2B).

### Disassembly of murine GFP-Mx1 nuclear bodies by 1,6-hexanediol

Rapid disassembly within 1-5 minutes of exposing cells to 1,6-hexanediol (3-5% w/v) in isotonic buffer is often used as a test of the liquid-like property of structures (6, 9, 27, 51-53). Hexanediol disrupts weak hydrophobic interactions that are required to preserve the structure of phase-separated condensates (51-53). The data in Fig. 3A represents a positive control for the rapid disassembly effect of hexanediol on cytoplasmic condensates of GFP-MxA in Mich-2H6 cells. Figs. 3B and 3C show illustrative images of the disassembly of GFP-Mx1 nuclear structures when Mich-2H6 and Huh7 cells were exposed to hexanediol. Compared to the rapid disassembly of cytoplasmic human GFP-MxA condensates (in less than 1 min; Fig. 3A), the hexanediol-induced disassembly of nuclear murine GFP-Mx1 is relatively slower (in 4-10 min; Figs. 3A and 3B). Overall, these hexanediol-triggered disassembly data provide evidence for GFP-Mx1 nuclear bodies to be phase-separated liquid-liquid (LLPS) condensates (9, 27, 51-53).

### Disassembly of GFP-Mx1 nuclear bodies by hypotonicity and reassembly by isotonicity

We and others reported previously that membraneless phase-separated condensates of GFP-MxA in the cytoplasm, or IL-6-induced cytoplasmic and nuclear GFP-STAT3 condensates, or influenza A virus (FLUAV) maturation structures in the cytoplasm were rapidly disassembled (within 1-3 minutes) in cells swollen following exposure to hypotonic buffer (9, 21, 27, 49). We further showed that a return of such cells to isotonic buffer caused rapid reassembly of MxA but in new structures different from the original ones (9, 27, 49). We have used the hypotonic disassembly-isotonic reassembly test to investigate whether GFP-Mx1 nuclear bodies might also represent such condensates.

As a negative control, Supplemental Fig. S1 shows that hypotonic-isotonic cycling had no effect on the dispersed N1-GFP protein expressed by itself in Huh7 cells. As another negative control, Fig. 5A in ref. 49 showed that dispersed cytoplasmic GFP-tagged STAT3 expressed in Huh7 cells also showed no change when cycled through hypotonic and isotonic buffers (in the absence of the addition of IL-6). In contrast, as a positive control, the data in Fig. 4A and Movie S1, show the rapid disassembly of cytoplasmic GFP-MxA condensates in Mich-2H6 cells. Moreover, Fig. 4A and Movie S1 also show the rapid reassembly of GFP-MxA in new condensates upon returning the cultures to isotonicity. Figs. 4B and 4C show illustrative images revealing the disassembly and reassembly of GFP-MuMx1 nuclear bodies following exposure of respective cells to hypotonic buffer and then return to isotonic buffer in both Mich2-H6 and Huh7 cells. These data further support the inference that GFP-MuMx1 nuclear bodies represent phase-separated condensates.

### FRAP assays on murine GFP-Mx1 nuclear bodies

Live-cell assays quantitating the speed of fluorescence recovery after photobleaching (FRAP) provide insight about the internal miscibility of GFP-tagged proteins within a condensate (1-5, 7, 9); rapid recovery indicates a more liquid interior, slower recovery suggests a more gel-like condensate (1-5, 7, 9). Thus, FRAP assays were carried out to investigate the mobility of GFP-Mx1 within the nuclear bodies. Fig. 5A and Movie S2 illustrate one example of this assay showing the photobleaching of the top and bottom parts of an elongated GFP-MuMx1 double nuclear structure followed by live-cell monitoring of recovery over the next 3 minutes. Fig. 5B compiles together several such assays (n =10). Taken together, the data show a slow recovery after photobleaching (mobile fraction = 18%), consistent with a gel-like consistency of the condensate (7, 9). This observation is reminiscent of the low mobility of GFP-HuMxA within cytoplasmic condensates (mobile fraction = 24%) (27). Thus, both GFP-HuMxA cytoplasmic condensates and GFP-MuMx1 nuclear condensates appear to have a gel-like internal consistency. As a positive control we have confirmed >70% mobile fraction in a FRAP assay when photobleaching soluble cytoplasmic GFP-STAT3 (27).

### Murine GFP-Mx1 is associated with a novel cytoplasmic giantin-based intermediate filament meshwork

Previous investigators have emphasized that murine Mx1 was exclusively observed in the nucleus (23-25, 29-31, 39-40). In the present experiments, we unexpectedly discovered that murine GFP-Mx1 expressed in Huh7 cells formed extensive cytoplasmic filamentous meshwork and often times were in cytoplasmic bodies in 20-30% of the transfected cells. Fig. 6A shows examples of such cells. Fig. 6A, left panel, shows not only a cell with extensive filamentous GFP-MuMx1 but also an adjacent cell which contains only nuclear bodies. Fig. 6A, right panel, shows a cell with not only cytoplasmic filaments but also cytoplasmic bodies. The cytoplasmic appearance of GFP-MuMx1 was not a result of “over-expression” in that often cells with only the cytoplasmic GFP-MuMx1 meshwork displayed weaker fluorescence intensity compared to cells with nuclear bodies. As controls we have verified that Huh7 cells expressing only the GFP protein or expressing GFP-STAT3 (a soluble protein) do not show any cytoplasmic or nuclear structures (Supplemental Fig. S1 and Fig. 5A in ref. 49).

The GFP-MuMx1 filaments in Huh7 cells were identified to be novel giantin-based intermediate filaments using giantin and vimentin as markers for this cytoskeleton network. We have previously reported that the elongated protein giantin was observed in Huh7 cells not only in the Golgi elements but also in intermediate filaments (27). As further verification, two different anti-giantin pAbs revealed this meshwork in Huh7 cells but not in Cos7 cells (Supplemental Fig. S2), a relevant giantin peptide but not an irrelevant STAT2 peptide competed off this immunofluorescence (Supplemental Fig. S3), and both giantin and vimentin colocalized in these filaments (Supplemental Fig. S4). As an additional feature, in cultures of Huh7 cells, the giantin-containing intermediate filaments extended between adjacent cells, perhaps along intercellular nanotubes (Fig. S5).

Fig. 6B shows colocalization of GFP-MuMx1 with the giantin-based intermediate filaments. The line-scan data show clearly that GFP-Mx1 filaments were also giantin positive. In Figs. 7A and Fig. 7B we evaluated the colocalization of vimentin with GFP-MuMx1 filaments. It is apparent that portions of the GFP-MuMx1 cytoplasmic meshwork were vimentin positive. Taken together, the data in Fig. 6B, 7A and 7B (together with the supporting data in Figs. S2-S5) identify the cytoplasmic GFP-MuMx1 filamentous meshwork in Huh7 cells as corresponding to a novel giantin-based intermediate filament cytoskeleton. Interestingly, we have previously reported that human GFP-MxA condensates also associated with this giantin-based intermediate filament meshwork in the cytoplasm of Huh7 cells (9, 27).

FRAP assays confirmed that photobleaching of GFP-Mx1 in these cytoplasmic filaments led to long-lived bleaching with little detectable recovery consistent with these filaments representing a long-lived cytoskeletal structure (Fig. 8 and Movie S3). Moreover, murine GFP-Mx1 associated with cytoplasmic filaments was not disassembled by either 1,6 hexanediol or hypotonicity (data not shown).

### Antiviral activity of murine GFP-Mx1 against VSV

It has been previously reported that murine Mx1 did *not* have an antiviral activity towards VSV, a rhabdovirus which replicates exclusively in the cytoplasm (25, 29, 30). With the realization that a significant subset of Huh7 cells expressed GFP-MuMx1 in cytoplasmic structures (Fig. 6 and Fig. 7), and the ability to investigate an antiviral activity towards VSV at the single-cell level using immunofluorescence methods (27, 30), we investigated whether cells expressing cytoplasmic GFP-MuMx1 might also display an antiviral phenotype towards VSV. Huh7 cultures were first transiently transfected with the GFP-MuMx1 expression vector or a GFP-HuMxA vector as a positive control and the cultures challenged two days later with VSV at a multiplicity of infection > 10 pfu/cell. The cultures were fixed 4 hour later, and the cells were probed for expression of viral nucleocapsid (N) protein by immunofluorescence. Images were quantitated at the single cell level using Image J for the levels of expression of N protein in cells showing no transfection, cytoplasmic GFP-HuMxA, only nuclear GFP-MuMx1, or only cytoplasmic GFP-MuMx1 (Fig. 9). Fig. 9A, 9B and 9C show representative examples of respective cells, while Fig. 9D summarizes the relevant quantitation. Fig. 9A and 9D confirm the antiviral activity of HuMxA towards VSV. Fig. 9B and 9D confirm the previous observations of Haller and colleagues (25, 29, 30) that cells expressing MuMx1 exclusively in nuclear bodies show no antiviral effect towards VSV. Remarkably, the data in Fig. 9C and 9D reveal that Huh7 cells with cytoplasmic MuMx1 have a strong antiviral phenotype towards VSV.

## Discussion

The IFN-induced antiviral proteins human MxA and its orthologous murine Mx1 localize predominantly to different cellular compartments – the human protein is cytoplasmic while the murine protein is, until now, thought to be exclusively nuclear (23-25, 29). We previously showed that human MxA formed membraneless phase-separated condensates and filaments in the cytoplasm (9, 26, 27). In the present study we provide evidence that the nuclear bodies formed by murine Mx1 are also phase-separated condensates. Moreover, FRAP analyses showed that, as with GFP-HuMxA condensates (mobile fraction: ∼24%; ref. 27), GFP-MuMx1 nuclear condensates had a gel-like internal consistency (mobile fraction: ∼18%). Additionally, we discovered that murine GFP-MuMx1 also participated in the formation of cytoplasmic bodies and a meshwork of cytoplasmic intermediate filaments in a subset of Huh7 cells. The latter observations suggested novel functions of MuMx1 in the cytoplasm, Indeed, we observed that cells with cytoplasmic MuMx1 displayed an antiviral activity towards VSV, a virus which replicates exclusively in the cytoplasm and one which had been previously reported to be unaffected by murine Mx1 (23-25). Thus, taken together with our previous data (26, 27), the present study shows (a) that both human MxA and murine Mx1 give rise to phase-separated biomolecular condensates, albeit in different cellular compartments, (b) both human MxA and murine Mx1 exhibit association with cytoplasmic intermediate filaments, at least in Huh7 cells, and (c) both human MxA and murine Mx1 when cytoplasmic exhibit an antiviral activity towards VSV – a rhabdovirus which replicates and matures entirely in the cytoplasm (Table 1).

**Table 1.**
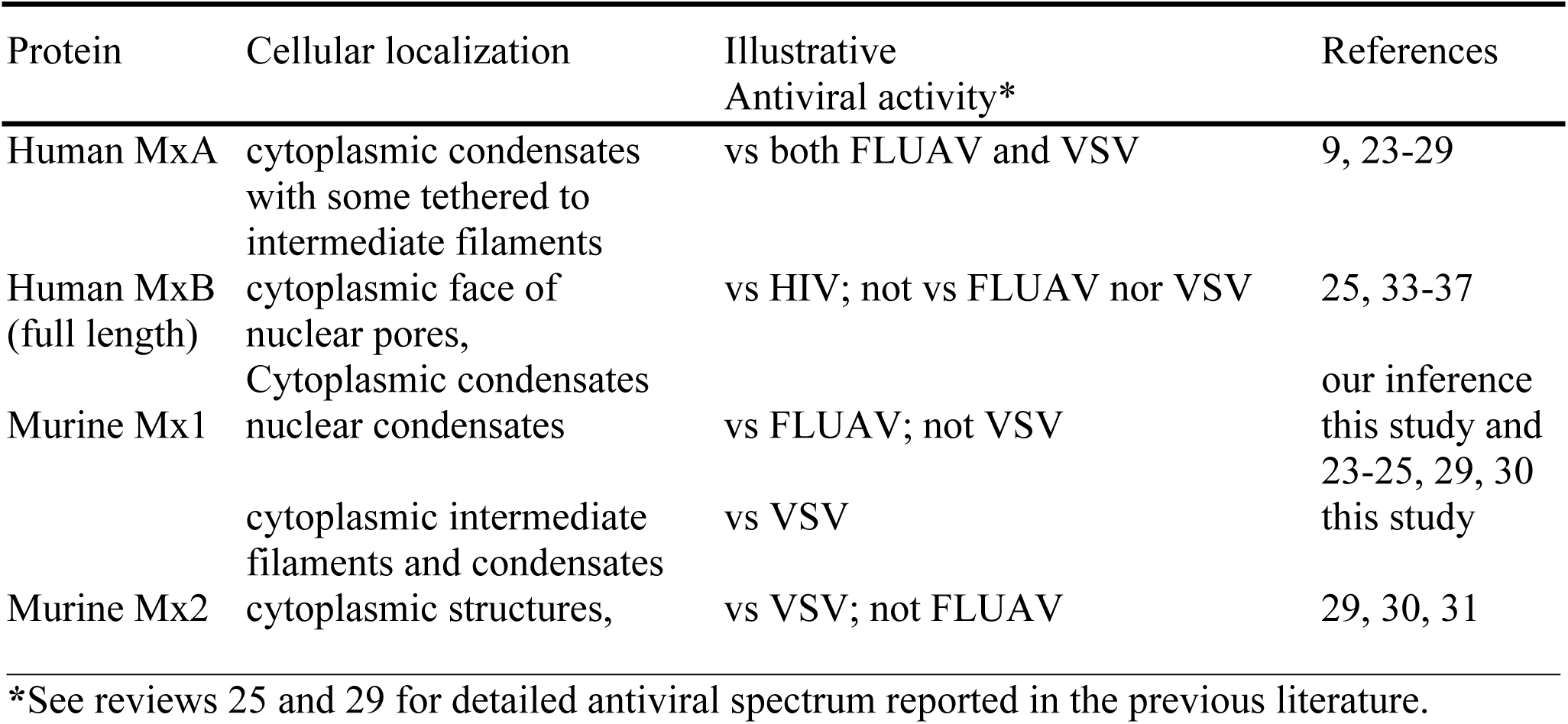
Properties of different human and murine Mx proteins.

The data showing the close relationship between MxB and nucleoporins on the cytoplasmic face of nuclear pores and the involvement of MxB in regulating cargo transit through the pore channel (33-37) allow us to now identify MxB structures at and near the nuclear pore to also represent phase-separated biomolecular condensates (Table 1; 51, 52). Previous research focused on characterizing the physical properties of various nucleoporins has already demonstrated these proteins to have the property of liquid-liquid phase separation (LLPS)(51, 52). Indeed, compounds such as hexanediol were used in cell biology to inhibit permeability through the pore channels through disruption of weak hydrophobic bonds between nucleoporin proteins in the pore condensates (51-53). Parenthetically, these observations eventually led to the more general use of disassembly by hexanediol as a test for phase-separated condensates (9, 53). Thus, the association of MxB with nuclear pore structures (33-37) taken together with the demonstration of nuclear pore structures to be phase-separated condensates (51, 52) points to the inference that MxB also associates with membraneless phase-separated condensates (see Figs. 2E and 2F in ref. 33 for examples).

Murine Mx1 typically shows antiviral activity towards influenza A virus (FLUAV) and other orthomyxoviruses which require a nuclear step in their replication, but not towards rhabdoviruses such as VSV which replicate entirely in the cytoplasm (23-25, 29) (Table 1). In contrast, functional murine Mx2 (isolated from feral mice) was observed to display an antiviral activity towards VSV but not FLUAV (29, 31). Commensurately, while murine Mx1 was observed to be in nuclear bodies, murine Mx2 was observed to be mainly in granular cytoplasmic structures (30, 31), which might perhaps also represent biomolecular condensates (Table 1)..

In an unexpected observation in the present study, we observed that a subset of GFP-MuMx1 expressing Huh7 cells contained cytoplasmic Mx1 structures, especially in association with novel giantin-based intermediate filaments. Remarkably, such cells displayed an antiviral activity towards VSV (Fig. 9C and 9D; Table 1). Such an antiviral activity was not observed in cells exclusively expressing GFP-MuMx1 in nuclear bodies (Fig. 9B and 9D). Thus, our data confirm the previous observation that cells expressing only murine Mx1 in the nucleus do not display an antiviral phenotype towards VSV (23-25), but now extend these observations to the discovery that murine GFP-Mx1 expressing Huh7 cells with cytoplasmic Mx1 expression show antiviral activity towards VSV. These data emphasize that the subcellular localization of respective Mx proteins affects the spectrum of Mx antiviral phenotypes observed. Curiously, it is notable that rat Mx1, which thus far has been reported to form only nuclear bodies, showed an antiviral phenotype against *both* FLUA and VSV (30, 38).

Extensive mutational studies of human MxA show that the GTPase activity is required for its antiviral activity, except that towards hepatitis B virus (23-25, 46, 54). However, data in the literature reveal that MxA mutants lacking GTase activity can still form cytoplasmic condensates (25, 54). However, mutations that cause dispersal of MxA in the cytoplasm (e.g. the D250N mutant) lacked antiviral activity (25, 46, 54). Moreover, a mutational analysis of rat Mx2 showed that mutants that formed “granular” cytoplasmic structures exhibited antiviral activity towards VSV, while those that were “diffuse” in the cytoplasm did not (38). These data suggest that condensate formation may be important for the antiviral activity of Mx proteins.

The hypotonicity-driven disassembly of Mx protein condensates in live cells (27, and present data) highlight an unusual aspect of Mx protein chemistry. The biochemical basis for this hypotonicity driven disassembly may reflect the effect of cytoplasmic “crowding” on higher-order protein structure or rapid changes due to hypotonicity-triggered post-translational modifications (4, 6, 9, 27). The reversal to isotonicity led to the reformation of both cytoplasmic GFP-HuMxA and nuclear GFP-MuMx1 condensates, albeit different from the original pre-hypotonicity condensates. These data suggest that the GFP-Mx proteins disperse completely under hypotonic conditions and then re-assemble upon reversal of cells to isotonicity to form new phase-separated condensates. It is noteworthy that virus infection often leads to the development of intracellular edema (55-58). Whether Mx proteins maintain their antiviral phenotype in the face of intracellular edema is not known.

Limitations of the present study include that in order to investigate single Mx species one at a time and without interference with endogenous Mx species we carried out transient transfection experiments using a vector for GFP-tagged Mx1 in heterologous human cells (Huh7 and Mich2-H6). These studies remain to be extended to investigations of GFP-MuMx1 in homologous murine cells as well as in MuIFN-treated murine cells. We have previously shown that, as with GFP-HuMxA transiently expressed in Huh7 cells, IFN-α-induced endogenous MxA also forms phase-separated cytoplasmic condensates (27).

To summarize, the present data taken together with those in the literature suggest that human MxA, its murine orthologs Mx1 and Mx2, as well as human MxB, likely have a common biophysical property – the ability to undergo liquid-liquid phase separation (LLPS) (Table 1). This results in the formation of different phase-separated biomolecular condensates [also called membraneless organelles (MLOs)] in different cellular compartments. Remarkably, the discovery of the antiviral activity of GFP-MuMx1 towards VSV further emphasizes the critical contribution of differences in subcellular localization in the biology of Mx proteins.

## Acknowledgements

We thank Drs. Joseph D. Etlinger (New York Medical College), Kenneth M. Lerea (New York Medical College) and Douglas S. Lyles (Wake Forest School of Medicine) for numerous helpful discussions. This research was supported, in part, by funds from New York Medical College, and by private funds of PBS. NYU Grossman School of Medicine Microscopy Laboratory is partially funded by NYU Cancer Center Support Grant NIH/NCI P30CA016087.

## Author contributions

PBS and AM designed the study, PBS carried out most of the experiments and imaging, HY carried out plasmid DNA preparations, HY and MFS carried out image quantitation and analyses, YD and FXL carried out confocal imaging and FRAP analyses, and PBS wrote the manuscript.

## Conflict of interest

All authors declare the absence of any conflict of interest

## Supplemental Movies

**Movie S1**. Time-lapse movie of the disassembly and reassembly of cytoplasmic human GFP-MxA condensates in Mich2-H6 cells. This movie corresponds to the experiment shown in Fig. 3, Panel A.

**Movie S2**. Time-lapse movie of the FRAP experiment on nuclear GFP-Mx1 condensates shown in Fig. 5A.

**Movie S3**. Time-lapse movie of the FRAP experiment on cytoplasmic filamentous GFP-Mx1 shown in Fig. 8.

## Legends to Supplemental Figures

**Fig. S1.**
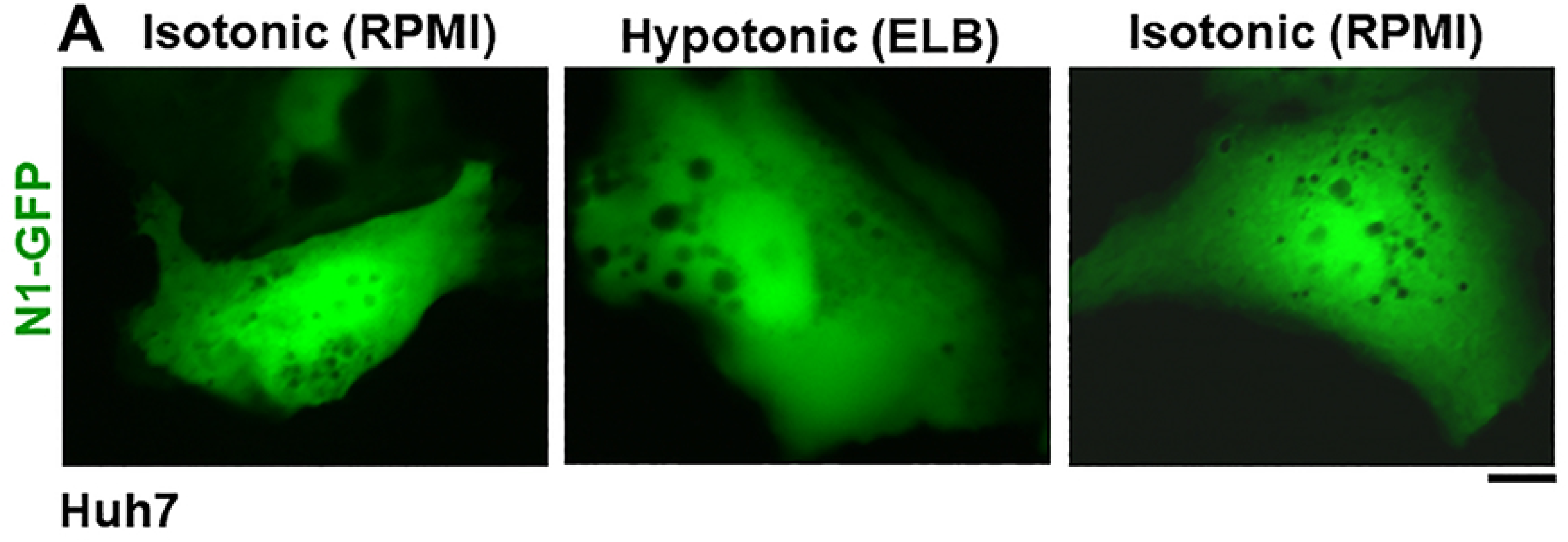
N1-GFP tag expressed in huh7 cells is not affected by hypotonic-isotonic cycling. Huh7 cells were transiently transfected with the N1-GFP vector as a negative control. One day later the live cells were first imaged in full medium (RPMI), and then the culture medium was changed to hypotonic ELB buffer (40 mOsm) followed by imaging at different times thereafter (9, 27). After 10-15 min, the culture medium was changed to isotonic RPMI live cell imaging continued for another 10-15 min. The figure illustrates representative cells at different time points during the experiment. Scale bar = 10 µm.

**Fig. S2.**
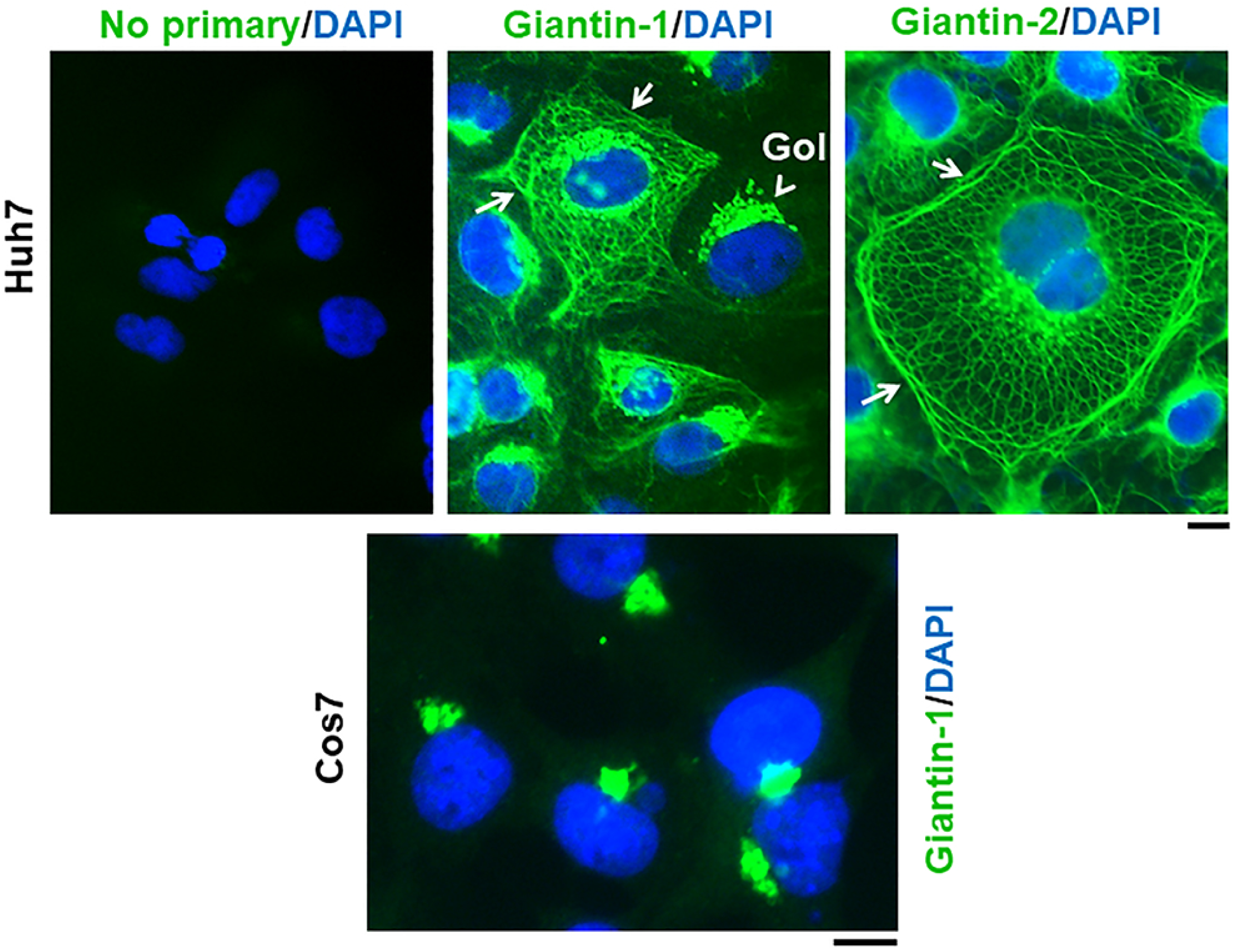
Giantin-based cytoplasmic filamentous meshwork in Huh7 hepatocytes. Huh7 and Cos7 cells grown in 35 mm culture dishes for 2 days were fixed using 4% paraformaldehyde, permeabilized using the 0.05% saponin/ 0.05% digitonin buffer (27) and immunostained with two different anti-giantin rabbit pAbs (giantin-1 was from Abcam, giantin-2 was from Biolegend) or processed without a primary antibody. All cultures were then exposed to AlexaFluor 488-tagged donkey anti-rabbit pAb and then DAPI staining. The cultures were imaged using three-color fluorescence. “Gol” points to the Golgi apparatus, white arrows point out the cytoplasmic filamentous meshwork in Huh7 cells. Scale bar = 10 µm.

**Fig. S3.**
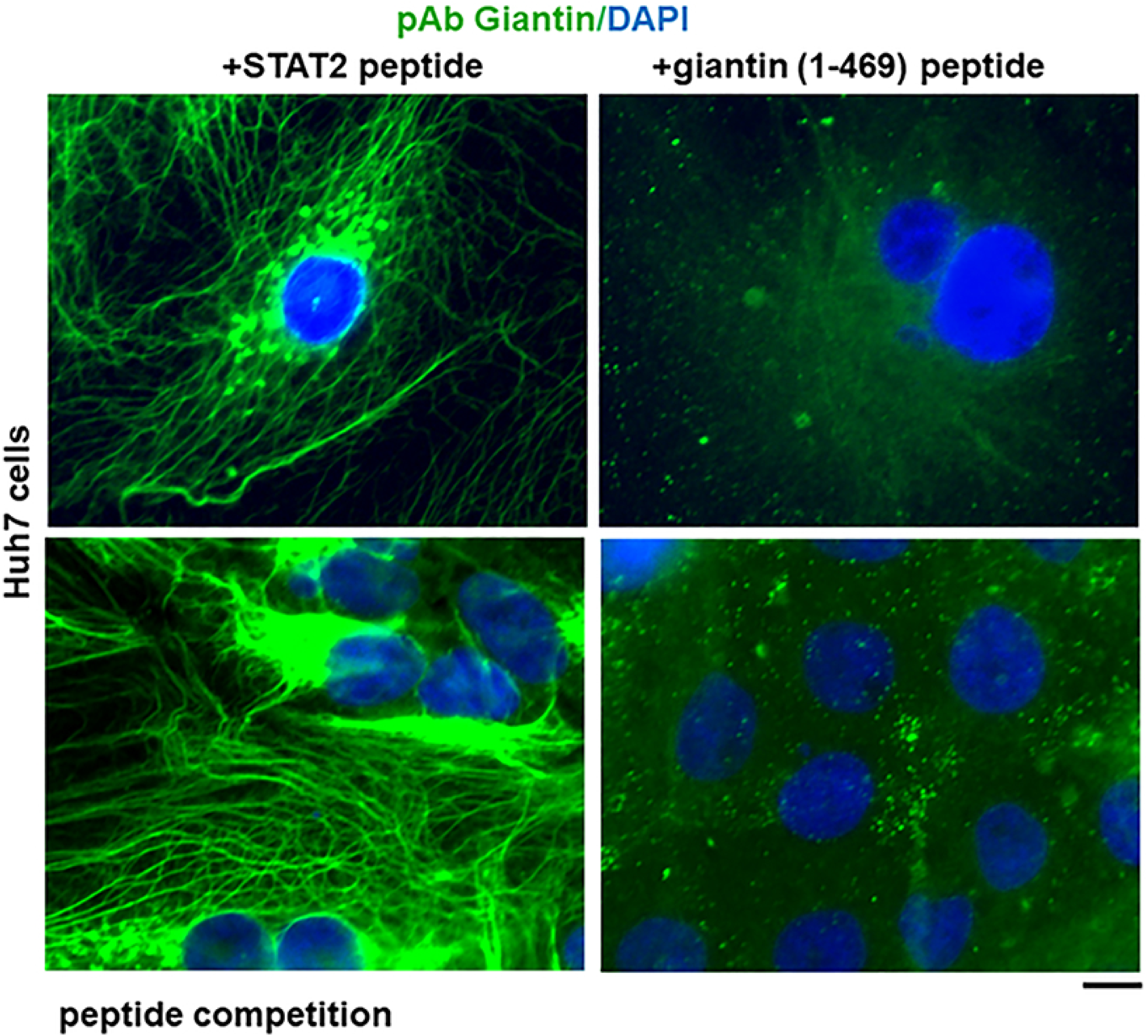
Peptide competition assay showing specificity of the giantin-based immunostaining of the cytoplasmic filamentous meshwork. Huh7 cells grown in 35 mm plates for 2 days were fixed using 4% paraformaldehyde, and permeabilized using the 0.05% saponin/0.05% digitonin buffer (27). Aliquots of anti-giantin pAb (Biolegend) were incubated with either STAT2 peptide or the relevant giantin (1-468) peptide for 30 min on ice and then used to carryout immunostaining of the respective cultures as in Fig. S2. The respective cultures were imaged at identical microscope settings. Scale bar = 10 µm.

**Fig. S4.**
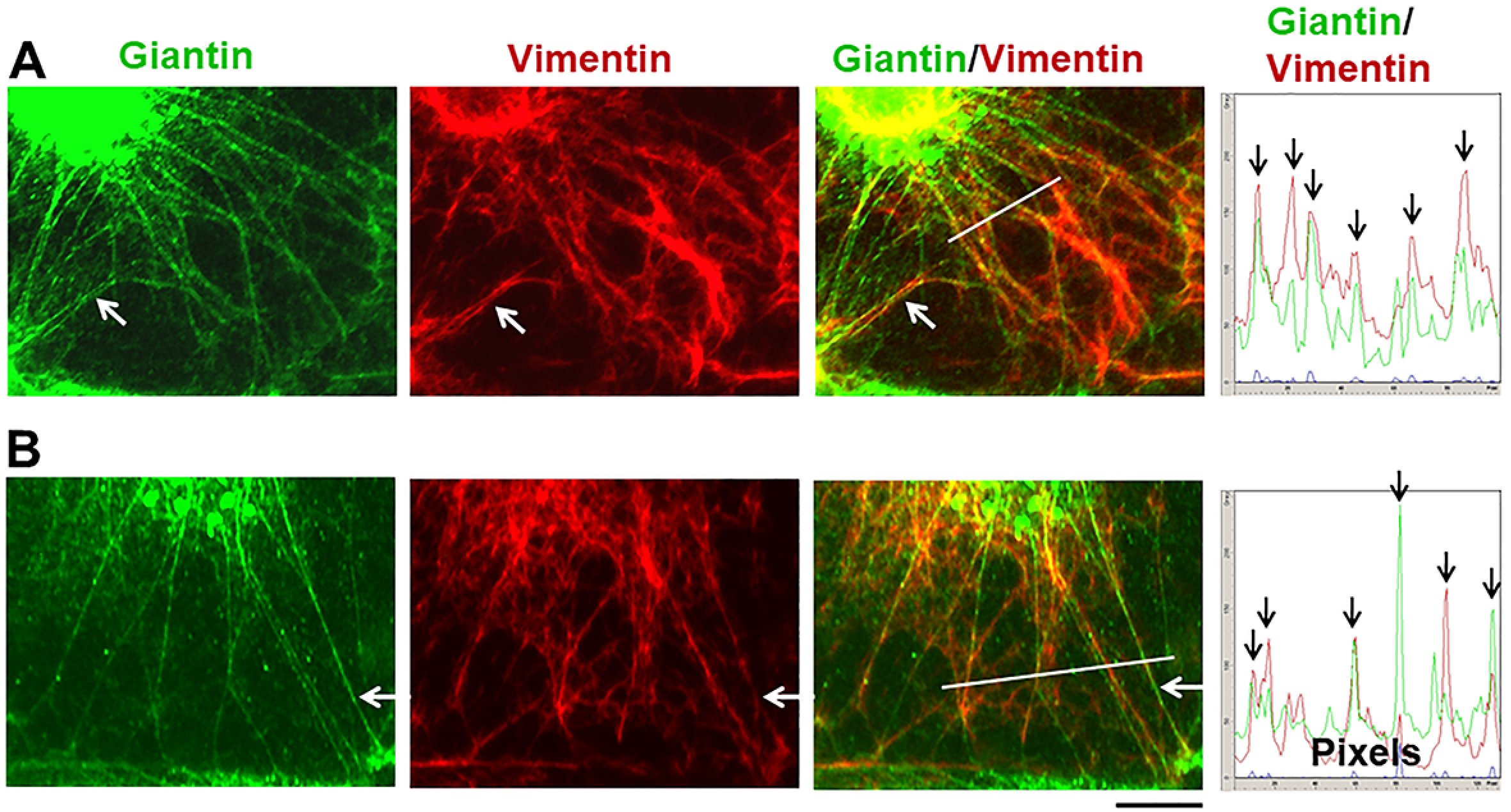
Identification of giantin-filaments in Huh7 cells as intermediate filaments. Huh7 cells grown in 35 mm plates for 2 days were fixed using 4% paraformaldehyde, permeabilized using the 0.05% saponin/0.05% digitonin buffer, and then subjected to double immunofluorescence analysis for vimentin and giantin in a sequential manner (27). Panels A and B illustrate two different cells. White arrows placed within the images point to filaments that are giantin and vimentin double positive. The plots on the right enumerate fluorescence intensity of red and green pixels along the white line in the respective merged images; black arrows point to giantin and vimentin double positive filaments in the respective scans. Scale bar = 10 µm.

**Fig. S5.**
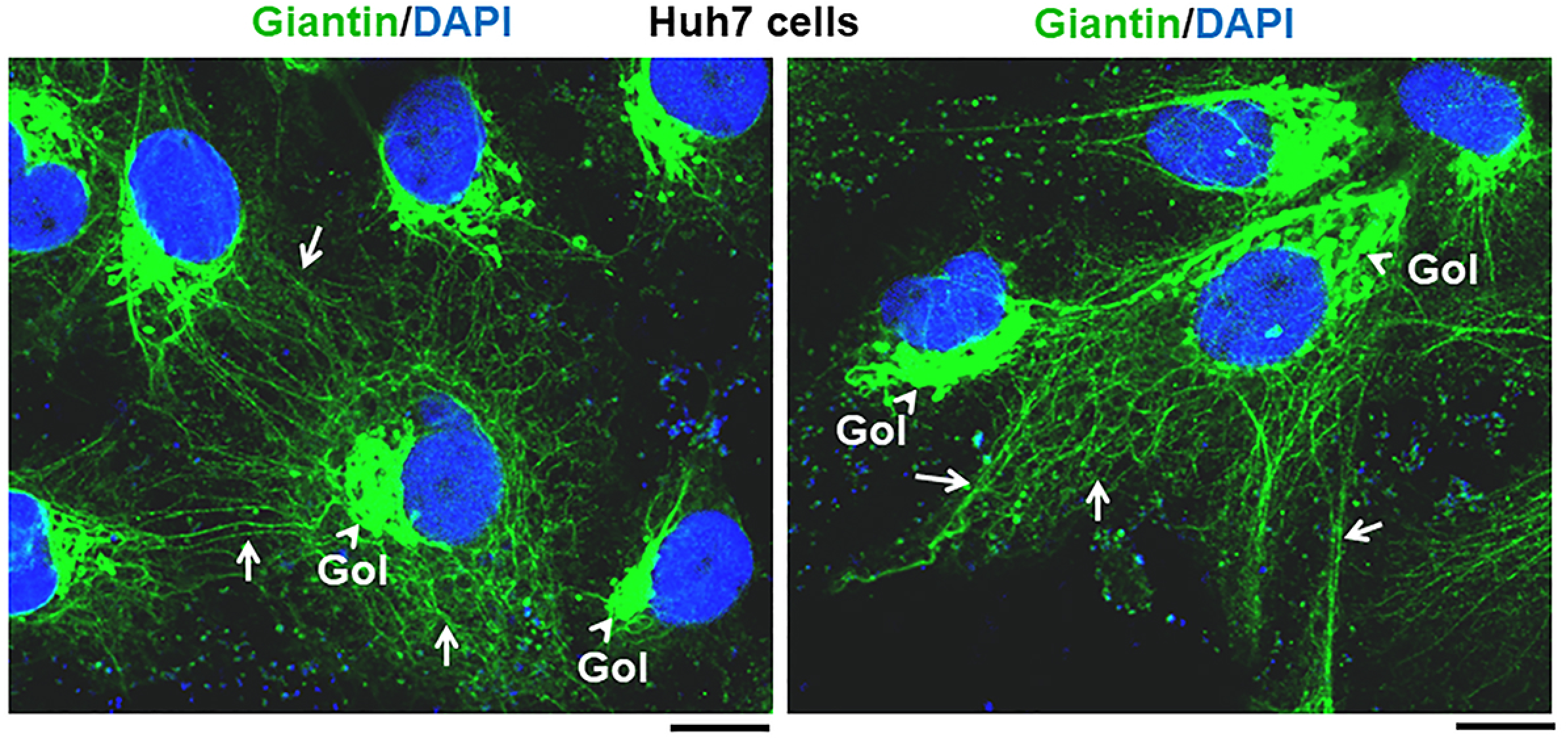
Giantin filaments extend between adjacent Huh7 cells. Huh7 cells grown in 35 mm plates for 2 days were fixed using 4% paraformaldehyde, permeabilized using the 0.05% saponin/0.05% digitonin buffer, and then subjected to immunofluorescence analysis for giantin followed by imaging using a high-resolution confocal microscope (27). Panels A and B illustrate two different regions of the culture. White arrows point to giantin-positive filaments many of which extend between adjacent cells (left panel). “Gol” points to the Golgi apparatus. Scale bar = 10 µm.

## Notes

### Competing Interest Statement

The authors have declared no competing interest.

## References

1. Mitrea, D.M. and Kriwacki, R.W. (2016) Phase separation in biology, functional organization of a higher order. Cell Commun. Signal 14, 1. doi: 10.1186/s12964-015-0125-7.

2. Banani, S.F., Lee, H.O., Hyman, A.A. and Rosen, M.K. (2017) Biomolecular condensates: organizers of cellular biochemistry. Nat Rev Mol Cell Biol. 18, 285–298.

3. Shin, Y. and Brangwynne, C.P. (2017) Liquid phase condensation in cell physiology and disease. Science 357(6357) pii: eaaf4382. doi: 10.1126/science.aaf4382.

4. Alberti, S. (2017) The wisdom of crowds: regulating cell function through condensed states of living matter. J Cell Sci. 130, 2789–2796.

5. Gomes, E. and Shorter. J. (2019) The molecular language of membraneless organelles. J Biol Chem 294, 7115–7127.

6. Alberti, S., Gladfelter, A. and Mittag, T. (2019) Considerations and challenges in studying liquid-liquid phase separation and biomolecular condensates. Cell 176, 419–434.

7. Brangwynne, C.P. (2018) Mapping local and global liquid phase behavior in living cells using photo-oligomerized seeds. Cell 175, 1467–1480.

8. Elbaum-Garfinkle, S. (2019) Matter over mind: liquid phase-separation and neurodegeneration. J Biol Chem 294, 7160–7168.

9. Sehgal, P. B., Westley, J., Lerea, K. M. DiSenso-Browne, S. and Etlinger, J. D. (2020) Biomolecular condensates in cell biology and virology: phase-separated membraneless organelles (MLOs). Analytical Biochem 597, 113691.

10. Reineke, L.C. and Lloyd, R.E. (2013) Diversion of stress granules and P-bodies during viral infection. Virology 436, 255–267.

11. Onomoto, K., Yoneyama, M., Fung, G., Kato H. and Fujita, T. (2014) Antiviral innate immunity and stress granule responses. Trends Immunol, 35, 420–428

12. Rozelle, D.K., Filone, C.M., Kedersha, N. and Connor, J.H. (2014) Activation of stress response pathways promotes formation of antiviral granules and restricts virus replication. Mol Cell Biol 34, 2003–2016.

13. Malinowska, M., Niedzwiedzka-Rystwej, P., Tokarz-Deptula, B. and Deptula, M. (2016) Stress granules (SG) and processing bodies (PB) in viral infections. Acta Biochima Polonica 63, 183–188.

14. McCormick, C. and Khapersky, D.A. (2017) Translation inhibition and stress granules in the antiviral immune response. Nat Rev Immunol 17, 647–660.

15. Heinrich, B.S., Maliga, Z., Stein, D.A., Hyman, A.A. and Whelan, S.P.J. (2018) Phase transitions drive the formation of vesicular stomatitis virus replication compartments. MBio 9, e02290–17. doi: 10.1128/mBio.02290-17.

16. Dinh, P.X., Beurs, L.K., Das, P.B., Panda, D., Das, A. and Pattnaik, A.K. (2013) Induction of stress granule-like structures in vesicular stomatitis virus-infected cells. J Virol 87, 372–383.

17. Nikolic, J., Bars, R.L., Lama, Z., Scrima, N., Lagaudriere-Gesbet, C., Gaudin, Y. and Blondel, D. (2017) Negri bodies are viral factories with properties of liquid organelles. Nat Commun 8, 58. doi: 10.1038/s41467-017-00102-9.

18. Hoenen, T., Shabman, R.S., Groseth, A., Herwig, A., Weber, M., Schudt, G., Dolnik, O., Basler, C.F., Becker, S. and Feldmann, H. (2012) Inclusion bodies are a site of ebolavirus replication. J. Virol 86, 11779–11788.

19. Zhou, Y., Su, J.M., Samuel, C.E. and Ma, D. (2019) Measles virus forms inclusion bodies with properties of liquid organelles. J Virol 93, e00948–19.

20. Guseva, S., Milles, G., Jensen, M. R., Salvi, N., Kleman, J. P., Maurin, D., Ruigrok, R. W. H. and Blackledge, M. (2020) Measles virus nucleo- and phosphoproteins form liquid-like phase-separated compartments that promote nucleocapsid assembly. Science Advances 6: eaaz7095 (April 1, 2020).

21. Alenquer, M., Vale-Costa, S., Sousa, A.L., Etibor, T.A., Ferreira, F. and Amorim. M.J. (2019) Influenza A virus ribonucleoproteins form liquid organelles at endoplasmic reticulum exit sites. Nature Comm 10, 1629. https://doi.org/10.1038/s41467-019-09549-4.

22. Peng, Q., Wang, L., Qin, Z., Wang, J., Zheng X., Wei, L., Zhang, X., Zhang, X. Liu, C., Li, Z., Wu, Y., Li, G., Yan, Q., Ma, J. (2020) Phase separation of Epstein-Barr virus EBNA2 and its coactivator EBNALP controls gene expression. J. Virol. DOI: 10.1128/JVI.01771-19

23. Haller, O. and Kochs, G. (2002). Interferon-induced mx proteins: dynamin-like GTPases with antiviral activity. Traffic 3, 710–717.

24. Haller, O., Staeheli, P., Kochs, G. (2007) Interferon-induced Mx proteins in antiviral host defense. Biochimie 89, 812–818.

25. Haller, O., Staeheli, P., Schwemmle, M. and Kochs, G. (2015) Mx GTPases: dynamin-like antiviral machines of innate immunity. Trends Microbiol 23, 154–163.

26. Davis, D., Yuan, H., Yang, Y.M., Liang, F.X. and Sehgal. P.B. (2018) Interferon-alpha-induced cytoplasmic MxA structures in hepatoma Huh7 and primary endothelial cells. Contemp Oncol (Pozn) 22, 86–94.

27. Davis, D., Yuan, H., Liang, F.X., Yang, Y.M., Westley, J., Petzold, C., Dancel-Manning, K., Deng, Y., Sall, J. and Sehgal, P.B. (2019) Human antiviral protein MxA forms novel metastable membraneless cytoplasmic condensates exhibiting rapid reversible tonicity-driven phase transitions. J Virol 93, e01014–19.

28. Kochs, G., Janzen, C., Hohenberg, H. and Haller, O. (2002) Antivirally active MxA protein sequesters La Crosse virus nucleocapsid protein into perinuclear complexes. Proc Natl Acad Sci USA 99, 3153–3158.

29. Verhelst, J., Hulpiau, P. and Saelens, X. (2013) Mx proteins: antiviral gatekeepers that restrain the uninvited. Microbiol. Mol. Biol. Rev. 77: 551–566.

30. Meier, E., Kunz, G, Haller, O. and Arnheiter, H. (1990) Activity of rat Mx proteins against a rhabdovirus. J. Virol. 64: 6263–6269.

31. Jin, H. K., Takada, A., Kon, Y., Haller, O., and Watanabe, T. (1999) Identification of the murine *Mx2* gene: interferon-induced expression of the Mx2 protein from the feral mouse gene confers resistance to vesicular stomatitis virus. J. Virol. 73: 4925–4930.

32. Busnadiego, I., Kane, M., Rihn, S., Preugchau, H. F., Hughes, J., Bianco-Melo, D., Strouvelle, V. P., Zang, T. M., Willett, B. J., Boutell, C., Bieniasz, P. D., Wilson, S. J. (2014) Host and viral determinants of Mx2 antiretroviral activity. J. Virol. 88: 7738–7752.

33. King, M. C., Raposo, G. and Lemmon, M. A. (2004) Inhibition of nuclear import and cell-cycle progression by mutated forms of the dynamin-like GTPase MxB. Proc. Natl. Acad. Sci. USA 101: 8957–8962.

34. Liu, Z., Pan, Q., Ding, S., Qian, J., Xu, F., Zhou, J., Cen, S., Guo, F. and Liang, C. (2013) The interferon-inducible MxB protein inhibits HIV-1 infection. Cell Host & Microbe 14: 398–410.

35. Goujon, C., Moncorge, O., Bauby, H., Doyle, T., Ward, C. C., Schaller, T., Hue, S., Barclay, W. S., Schulz, R. and Malim, M. H. (2013) Human Mx2 is an interferon-induced post-entry inhibitor of HIV-1 infection. Nature 502: 559–562.

36. Goujon, C., Moncorge, O., Bauby, H., Doyle, T., Barclay, W. S. and Malim, M. H. (2014) Transfer of the amino-terminal nuclear envelope targeting domain of human MX2 converts MX1 into an HIV-1 resistance factor. J. Virol. 88: 9017–9026.

37. Dicks, M. D. J., Belancor, G., Jimenez-Guardeno, J. M., Pessei-Vivares, L., Apolonia, L., Goujon, and Malim, M. H. (2018) Multiple components of the nuclear pore complex interact with the amino-terminus of MX2 to facilitate HIV-1 restriction. PLoS Pathog 14(11): e1007408.

38. Johannes, L., Kambadur, R., Lee-Hellmich, H., Hodgkinson, C. A., Arnheiter, H. and Meier, E. (1997) Antiviral determinants of rat Mx GTPases map to the carboxy-terminal half. J. Virol. 71: 9792–9795.

39. Engelhardt, O.G., Ullrich, E., Kochs. G. and Haller, O. (2001) Interferon-induced antiviral Mx1 GTPase is associated with components of the SUMO-1 system and promyelocytic leukemia protein nuclear bodies. Exptl Cell Res 271, 286–295.

40. Engelhardt, O.G., Sirma, H., Pandolfi, P.P. and Haller, O. (2004) Mx1 GTPase accumulates in distinct nuclear domains and inhibits influenza A virus in cells that lack promyelocytic leukemia protein nuclear bodies. J Gen Virol 85, 2315–2326.

41. Chelbi-Alix, M.K., Pelicano, L., Quignon, F., Koken, M.H., Venturini, L., Stadler, M., Pavlovic, J., Degos. L. and de The, H. (1995) Induction of the PML protein by interferons in normal and APL cells. Leukemia 9, 2027–2033.

42. Blight KJ, McKeating JA, Rice CM. 2002. Highly permissive cell lines for subgenomic and genomic hepatitis C virus RNA replication. J Virol 76:13001–13014.

43. Mackiewicz, J., Karczewaska-Dzionk, A., Laciack, M., Kapcinska, M., Wiznerowicz, M, Burkowski, T., Zakowska, M., Rose-John, S. and Mackiewicz, M. (2015) Whole cell therapeutic vaccine modified with huper-IL6 for combinational treatment of nonresected advanced melanoma. Medicine 94:e853.

44. Lee JE, Yang YM, Liang FX, Gough DJ, Levy DE, Sehgal PB. 2012. Nongenomic STAT5-dependent effects on Golgi apparatus and endoplasmic reticulum structure and function. Am J Physiol Cell Physiol 302:C804–C820.

45. Yuan H, Sehgal PB. 2016. MxA Is a Novel Regulator of Endosome-Associated Transcriptional Signaling by Bone Morphogenetic Proteins 4 and 9 (BMP4 and BMP9). PLoS One 11:e0166382.

46. Wisskirchen C, Ludersdorfer TH, Muller DA, Moritz E, Pavlovic J. 2011. Interferon-induced antiviral protein MxA interacts with the cellular RNA helicases UAP56 and URH49. J Biol Chem 286:34743–34751.

47. Nigg PE, Pavlovic J. 2015. Oligomerization and GTP-binding Requirements of MxA for Viral Target Recognition and Antiviral Activity against Influenza A Virus. J Biol Chem 290:29893–29906.

48. Xu, F., Mukhopadhyay, S. and Sehgal, P.B. (2007) Live cell imaging of interleukin-6-induced targeting of “transcription factor” STAT3 to sequestering endosomes in the cytoplasm. Am J Physiol Cell Physiol 293, C1374–1382.

49. Sehgal, P. B. (2019) Biomolecular condensates in cancer cell biology: interleukin-6-induced cytoplasmic and nuclear STAT3/PY-STAT3 condensates in hepatoma cells. Contemp Oncol (Pozn). 23, 16–22.

50. Carey BL, Ahmed M, Puckett S, Lyles DS. 2008. Early steps of the virus replication cycle are inhibited in prostate cancer cells resistant to oncolytic vesicular stomatitis virus. J Virol 82:12104–12115.

51. Ribbeck, K. and Gorlich. D. (2002) The permeability barrier of nuclear pore appears to operate via hydrophobic exclusion. EMBO Journal 21, 2664–2671.

52. Patel, S. S., Belmont, B. J., Santa, J. M. and Rexach, M. F. (2007) Natively unfolded nucleoporins gate protein diffusion across the nuclear pore complex. Cell 129, 83–96.

53. Kroschwald, S., Maharana, S., Mateju, D., Malinovska, L., Nuske, E., Poser, I, Richter. and Alberti, S (2015). Promiscuous interactions and protein disaggregases determine the material state of stress-inducible RNP granules. eLIFE 4, e06807.

54. Dick A, Graf L, Olal D, von der Malsburg A, Gao S, Kochs G, Daumke O. 2015. Role of nucleotide binding and GTPase domain dimerization in dynamin-like myxovirus resistance protein A for GTPase activation and antiviral activity. J Biol Chem 290:12779–12792.

55. Knipe DM. 1990. Virus-host-cell interactions, p 293–316. In Fields BN, Knipe DM (ed), Fields’ Virology, 2nd ed, vol 1. Raven Press.

56. Fernandez-Puentes C, Carrasco L. 1980. Viral infection permeabilizes mammalian cells to protein toxins. Cell 20:769–75.

57. Carrasco L. 1978. Membrane leakiness after viral infection and a new approach to the development of antiviral agents. Nature 272:694–699.

58. Carrasco L, Smith AE. 1976. Sodium ions and the shut-off of host cell protein synthesis by picornaviruses. Nature 264:807–809.

